# Pre-existing interferon gamma conditions the lung to mediate early control of SARS-CoV-2

**DOI:** 10.1101/2023.07.15.549135

**Authors:** Kerry L. Hilligan, Sivaranjani Namasivayam, Chad S. Clancy, Paul J. Baker, Samuel I. Old, Victoria Peluf, Eduardo P. Amaral, Sandra D. Oland, Danielle O’Mard, Julie Laux, Melanie Cohen, Nicole L. Garza, Bernard A. P. Lafont, Reed F. Johnson, Carl G. Feng, Dragana Jankovic, Olivier Lamiable, Katrin D. Mayer-Barber, Alan Sher

**Affiliations:** Immunobiology Section, Laboratory of Parasitic Diseases, National Institute of Allergy and Infectious Diseases, National Institutes of Health, Bethesda, MD 20892, USA; Malaghan Institute of Medical Research, Wellington 6012, New Zealand; Rocky Mountain Veterinary Branch, National Institute of Allergy and Infectious Diseases, National Institutes of Health, Hamilton, MT 59840, USA; Inflammation and Innate Immunity Unit, Laboratory of Clinical Immunology and Microbiology, National Institute of Allergy and Infectious Diseases, National Institutes of Health, Bethesda, MD 20892, USA; Immunoparasitology Unit, Laboratory of Parasitic Diseases, National Institute of Allergy and Infectious Diseases, National Institutes of Health, Bethesda, MD 20892, USA; Flow Cytometry Section, Research Technologies Branch, National Institute of Allergy and Infectious Diseases, National Institutes of Health, Bethesda, MD 20892, USA; SARS-CoV2-Virology Core, Laboratory of Viral Diseases, National Institute of Allergy and Infectious Diseases, National Institutes of Health, Bethesda, MD 20892, USA; Immunology and Host Defense Group, School of Medical Sciences, Faculty of Medicine and Health, The University of Sydney, NSW 2006, Australia; Centenary Institute, The University of Sydney, NSW 2050, Australia

## Abstract

Interferons (IFNs) are critical for anti-viral host defence. Type-1 and type-3 IFNs are typically associated with early control of viral replication and promotion of inflammatory immune responses; however, less is known about the role of IFNγ in anti-viral immunity, particularly in the context of SARS-CoV-2. We have previously observed that lung infection with attenuated bacteria *Mycobacterium bovis* BCG achieved though intravenous (*iv*) administration provides strong protection against SARS-CoV-2 (SCV2) infection and disease in two mouse models. Assessment of the pulmonary cytokine milieu revealed that *iv* BCG induces a robust IFNγ response and low levels of IFNβ. Here we examined the role of ongoing IFNγ responses due to pre-established bacterial infection on SCV2 disease outcomes in two murine models. We report that IFNγ is required for *iv* BCG induced reduction in pulmonary viral loads and that this outcome is dependent on IFNγ receptor expression by non-hematopoietic cells. Further analysis revealed that BCG infection promotes the upregulation of interferon-stimulated genes (ISGs) with reported anti-viral activity by pneumocytes and bronchial epithelial cells in an IFNγ-dependent manner, suggesting a possible mechanism for the observed protection. Finally, we confirmed the importance of IFNγ in these anti-viral effects by demonstrating that the recombinant cytokine itself provides strong protection against SCV2 challenge when administered intranasally. Together, our data show that a pre-established IFNγ response within the lung is protective against SCV2 infection, suggesting that concurrent or recent infections that drive IFNγ may limit the pathogenesis of SCV2 and supporting possible prophylactic uses of IFNγ in COVID-19 management.

## Introduction

COVID-19 is a pulmonary disease caused by SARS-CoV-2 (SCV2) which infects lung epithelial cells via the membrane protein angiotensin-converting enzyme 2 (ACE2) (*1, 2*). ACE2 is highly expressed by pneumocytes and particular subsets of ciliated bronchial cells (*3, 4*), thus making these cell types the primary target for SCV2 infection (*5*). Interferons (IFNs) play a central role in anti-viral immunity, through induction of host defense elements that constrain viral invasion, replication, and release in target epithelial cells (*6, 7*), as well as by facilitating immune cell activation and recruitment (*8*). Type-1 IFNs (IFN-I, including IFNα and IFNβ) and type-3 IFNs (IFNλ) are typically associated with responses against viruses and have been identified as mediators of host defense against SCV2 (*9–14*). However, the role of type-2 IFN (IFNγ) during viral infection and in particular, SCV2 infection, is less clear.

IFNγ is a key mediator of immunity to intracellular microbes and is strongly induced upon bacterial infection. Natural killer (NK) and innate lymphoid cells contribute innate sources of IFNγ, whereas CD4+ Th1 cells and CD8+ T cells are major producers of this cytokine later in infection. An important function of IFNγ is to arm myeloid cells with microbiocidal properties such as induction of nitric oxide synthase (NOS)-2, which can also inhibit some viruses (*15*). In addition, IFNγ broadly induces a suite of interferon-stimulated genes (ISGs), many of which are also induced by type-1 and type-3 IFNs and have been reported to also possess anti-viral activity (*16*). While IFNγ is not generally required for host resistance to a pulmonary viral infection (*17–21*), recombinant (r)IFNγ treatment has been shown to confer protection against certain viral pathogens in animal model studies (*22–26*). In COVID-19 patients, including immunocompromised individuals, rIFNγ treatment was shown to be well tolerated (*27, 28*) and in one study was reported to reduce the time to hospital discharge (*29*).

Pre-clinical studies and clinical trials assessing the efficacy of therapeutics that act through IFNs have shown that the timing of treatment relative to viral exposure is crucial (*25, 30–32*). Successful regimens usually involve treatment before or just after viral exposure limiting their potential therapeutic use. However, these observations do raise some interesting questions concerning host susceptibility to SCV2 infection in the context of an ongoing IFN response to concurrent or recent infection. Indeed, we and others have previously observed that lung infection with *Mycobacterium bovis* BCG achieved through intravenous (*iv*) administration provides protection against SCV2 in mouse and hamster models (*33–35*). Likewise, aerosol *Mycobacterium tuberculosis* infection has been shown to be associated with lower SCV2 viral burdens and improved survival in mice (*36, 37*).

Here we explored the mechanisms by which concurrent bacterial infection protects against SCV2 and show that IFNγ is an essential mediator of BCG conferred protection *in vivo*. IFNγ was found to act on epithelial cells to limit SCV2 infection and/or replication, possibly through the induction of anti-viral proteins. Intranasal administration of the recombinant cytokine prior to viral challenge also elicited strong protection against SCV2, recapitulating our observations in bacterial co-infection models. Together these observations support a role for pre-existing IFNγ responses in mediating early control of SCV2.

## Results

### *iv* BCG alters the pulmonary cellular landscape and promotes a strong IFNγ signature

We and others have observed that intravenous (*iv*), but not subcutaneous (*sc*), administration of BCG protects against SCV2 in mice and hamsters (*33–35*), thus providing a platform to dissect mechanisms of host resistance to SCV2. To gain a deeper understanding of *iv* BCG mediated protection against SCV2, we set up a new series of experiments where BCG or PBS was administered *iv* to wildtype (WT) B6 mice 40-45 days before SCV2 infection, at which time there is ∼10^5^ BCG CFU present in the lung (*33*). In these experiments, we utilized a beta variant (B.1.351) of the virus carrying a N501Y mutation that can transiently infect WT mice through binding of the endogenous murine ACE2 receptor (*38–42*) (**Fig 1A**). As we had previously observed with an alpha variant of SCV2 (*33*), *iv* BCG significantly reduced lung viral burden at 3 days after challenge with a beta variant (**Fig 1B**). To identify correlates of protection, we performed single-cell RNA sequencing (scRNASeq) on lung cells isolated from the same set of animals. Seurat clustering revealed 28 distinct cell types encompassing epithelial, endothelial, stromal, myeloid and lymphoid lineages (**Fig 1C, FigS1A and Table S1**). When comparing the abundance of different clusters between control (PBS) and *iv* BCG animals, we found CD4+ T cells and CD8+ T cells to be enriched after *iv* BCG and fibroblast subsets to be comparatively higher in controls (**Fig 1D**). This is in line with flow cytometry data showing significantly higher numbers of T lymphocytes present in the lung tissue of *iv* BCG mice compared to controls, which is also apparent prior to SCV2 challenge (**FigS1B-C**).

**Figure 1:**
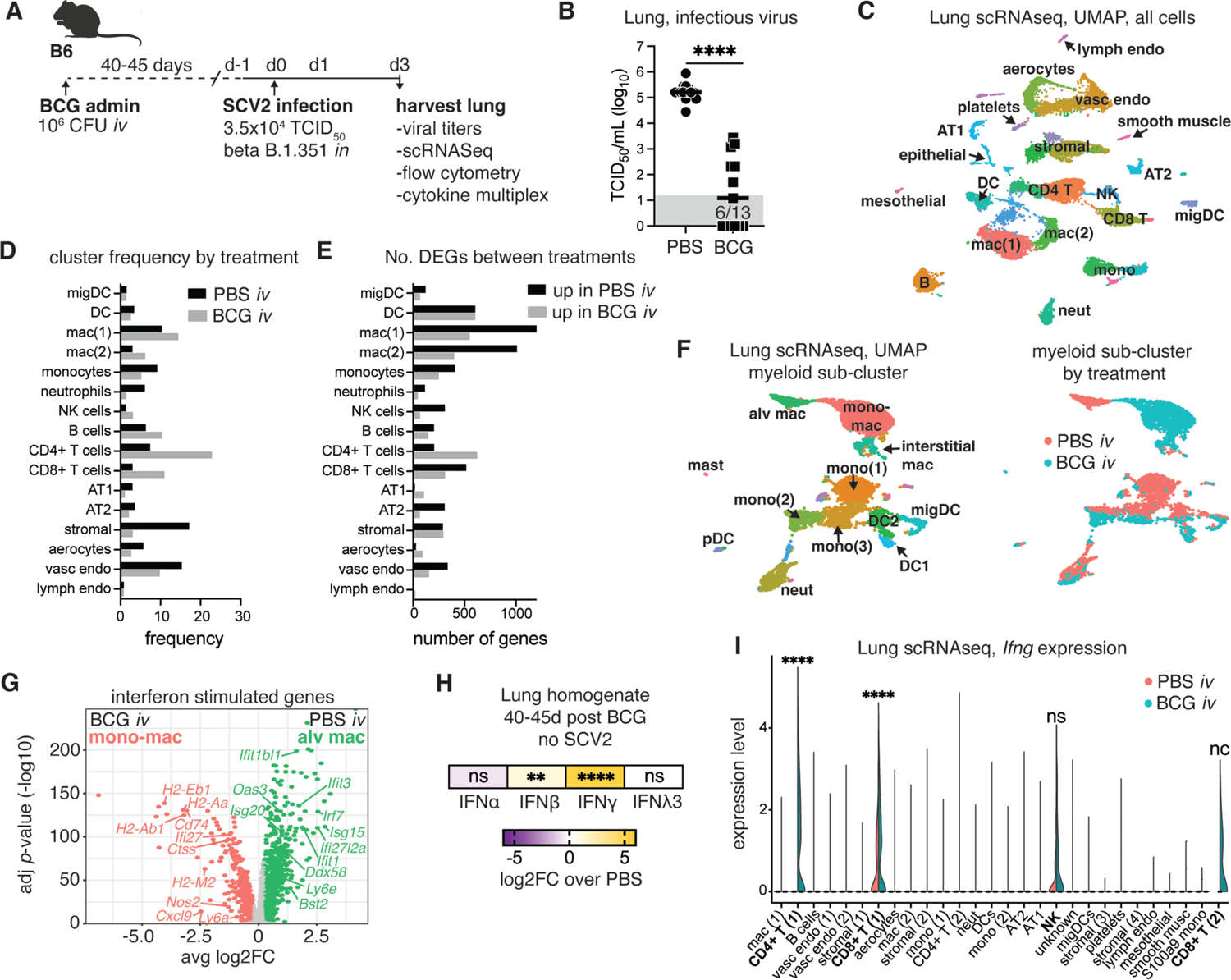
*iv* BCG skews the pulmonary cytokine landscape towards IFNγ production. B6 mice were inoculated with BCG or PBS *iv* 40-45 days prior to intranasal challenge with SCV2 B.1.351. Lungs were harvested 3 days after viral challenge. (**A**) Schematic of experimental protocol. (**B**) Viral titers in lung homogenate as measured by TCID_50_ assay. Data are pooled from 2 independent experiments with 5-9 mice/group. Statistical significance was assessed by Mann-Whitney test. \*\*\*\**p*<0.0001. Gray box shows values below limit of detection. (**C**) UMAP representation of scRNAseq data of whole lung isolated from BCG and PBS animals challenged with SCV2 B.1.351. Clustering resolution: 0.6. Cell clusters were manually annotated. Cluster specific annotation can be found in FigS1 and gene sets found in Supplementary Table 1. (**D**) Frequency of cells in each cluster separated by experimental condition. (**E**) Number of differentially expressed genes (FC>2.5, *p*<0.05) between PBS and BCG groups for each cluster. Genes more highly expressed in the PBS sample are shown in the black bars. Genes more highly expressed in the BCG sample are shown in the gray bars. Gene lists with their associated FC and *p*-values can be found in Supplementary Table 2. (**F**) UMAP representation of re-clustered myeloid cells for PBS or BCG treated animals (left and UMAP colored by treatment group (right). Clustering resolution: 0.4. Cell clusters were manually annotated. Cluster specific gene sets can be found in Supplementary Table 3. (**G**) Volcano plot shows DEGs with annotated ISGs between resident AM and monocyte-derived macrophage clusters. The full gene list and their associated FC and *p*-values can be found in Supplementary Table 4. (**H**) Fold change in interferon protein levels in lung homogenate between PBS and BCG treated mice. Data are pooled from 3 independent experiments with 4-5 mice/group. Statistical significance was assessed by unpaired t-tests. Not significant (ns) *p*>0.05; **, *p*<0.01; \*\*\*\**p*<0.0001. (**I**) Violin plots show *Ifng* expression across UMAP clusters from Fig 1C separated by experimental condition. Statistical significance was assessed by Wilcoxon Rank Sum test with Bonferroni correction. Not significant (ns) *p*>0.05; \*\*\*\**p*<0.0001. Not calculated (nc) due to cluster only containing cells from one treatment group.

We next performed differential expression analysis of scRNASeq data which showed that T cell and myeloid cell clusters, in particular macrophage populations mac(1), mac(2) and dendritic cells (DC), had the highest number of differentially expressed genes (DEGs) between PBS and BCG treated mice (**Fig 1E and Table S2**). Sub-clustering of the myeloid compartment identified 16 clusters encompassing macrophages, monocytes, dendritic cells, neutrophils and mast cells, many of which showed prominent condition-specific clustering (**Fig 1F, FigS1D and Table S3**). Notably, resident alveolar macrophages were found exclusively in samples from control animals, whereas *iv* BCG administration was associated with monocyte-derived macrophages (**Fig 1F**). In addition to genes related to cell ontogeny (**FigS1E**), DEGs between these two macrophage populations were related to inflammatory responses, anti-viral signatures and interferon signaling, with the alveolar macrophages from control animals expressing an IFN-I signature (*Isg15*, *Oas3*, *Ifitm3*) consistent with the high viral titers recovered from the lungs. In contrast, the monocyte-derived macrophages from BCG treated mice expressed robust glycolytic, antigen presentation and IFNγ signatures including high levels of *Nos2*, *Cxcl9* and numerous MHCII related genes (**Fig 1G, FigS1E and Table S4**). This observation is in line with cytokine multiplex and ELISA data from SCV2 naïve animals showing IFNα and IFNλ3 levels did not differ between PBS and BCG groups, whereas IFNβ and IFNγ were significantly higher in samples from BCG-treated animals compared to controls, with the most striking increase seen in the IFNγ levels (**Fig 1H**).

To evaluate the sources of IFNγ after *iv* BCG, we mapped *Ifng* transcripts against the UMAP projection of the single-cell RNAseq clustering. This revealed that *Ifng* is predominantly expressed by CD4+ and CD8+ T lymphocytes as well as NK cells in *iv* BCG mice. Only low-level expression was observed in control mice which was largely restricted to CD8+ T cells and NK cells (**Fig 1I**). While these data are from 3 days after SCV2 challenge, they are consistent with flow cytometric analysis showing increased CD4+ T cell expression of the Th1 master transcription factor Tbet 42 days after *iv* BCG with no SCV2 challenge (**FigS1F**).

Together these data show that *iv* BCG induces a T and NK cell driven IFNγ response in the lung, which is also apparent prior to SCV2 challenge. Importantly, elevated IFNγ responses levels and Tbet+ Th1 cells were only observed after *iv* administration and not in SCV2-susceptible animals administered BCG by the *sc* route (**FigS1F**) (*33*). This route dependent induction of Th1 cells producing IFNγ is likely due the fact that only *iv* administration establishes a substantial bacterial infection of the lung (*33*). We therefore hypothesized that a bacteria prompted IFNγ response within the pulmonary micro-environment prior to SCV2 exposure may be involved in the protection conferred by *iv* BCG.

### IFNγ is required for *iv* BCG conferred protection against SCV2

We next evaluated whether IFNγ receptor signaling and T cells, the major source of IFNγ after *iv* BCG, are required for protection against SCV2 in response to *iv* BCG. To this end, BCG was administered *iv* to WT B6, *Ifngr1*^-/-^ or *Tcra*^-/-^ mice 40-45 days before intranasal SCV2 challenge. At three days post SCV2 challenge, viral loads were assessed in the lung by tissue culture infectious dose-50 (TCID_50_) assay (**Fig 2A**). Strikingly, in the absence of the IFNγ receptor, prior *iv* BCG infection failed to reduce viral loads as seen in WT B6 mice. Similarly, BCG protection was diminished in T cell deficient animals, although some reduction in viral load was still noted, potentially due to residual IFNγ produced by NK cells in these animals (**Fig 2B**). Mice with deficiencies in IFNγ signaling and/or T cells are highly susceptible to BCG (*43, 44*) and consistent with this, the colony forming units (CFU) recovered from the lungs of *Ifngr1*^-/-^ or *Tcra*^-/-^ animals were significantly higher than WT B6 controls (**FigS2**). To control for this difference in CFU burden, we adopted an alternative approach where WT B6 mice were inoculated *iv* with BCG and IFNγ was subsequently neutralized through intraperitoneal administration of an anti-IFNγ antibody starting just 1 day prior to SCV2 challenge (**Fig 2C**). Under these experimental conditions, no increase in BCG CFU was observed in the lungs after IFNγ neutralization (**FigS2**), yet SCV2 viral loads were strikingly higher than in isotype control animals inoculated *iv* with BCG (**Fig 2D**). As some residual protection was observed in *iv* BCG mice after IFNγ neutralization, we also blocked the IFN-I receptor (IFNAR) immediately prior to SCV2 challenge to determine whether the low levels of IFNβ detected after *iv* BCG could also be contributing to the anti-SCV2 response. This treatment failed to further reduce the protection achieved by administration of the anti-IFNγ antibody alone (**Fig 2D**). Together with the data showing complete loss of protection in *Ifngr1*^-/-^ mice, this result implicated IFNγ as the dominant mediator of *iv* BCG conferred protection against SCV2.

**Figure 2:**
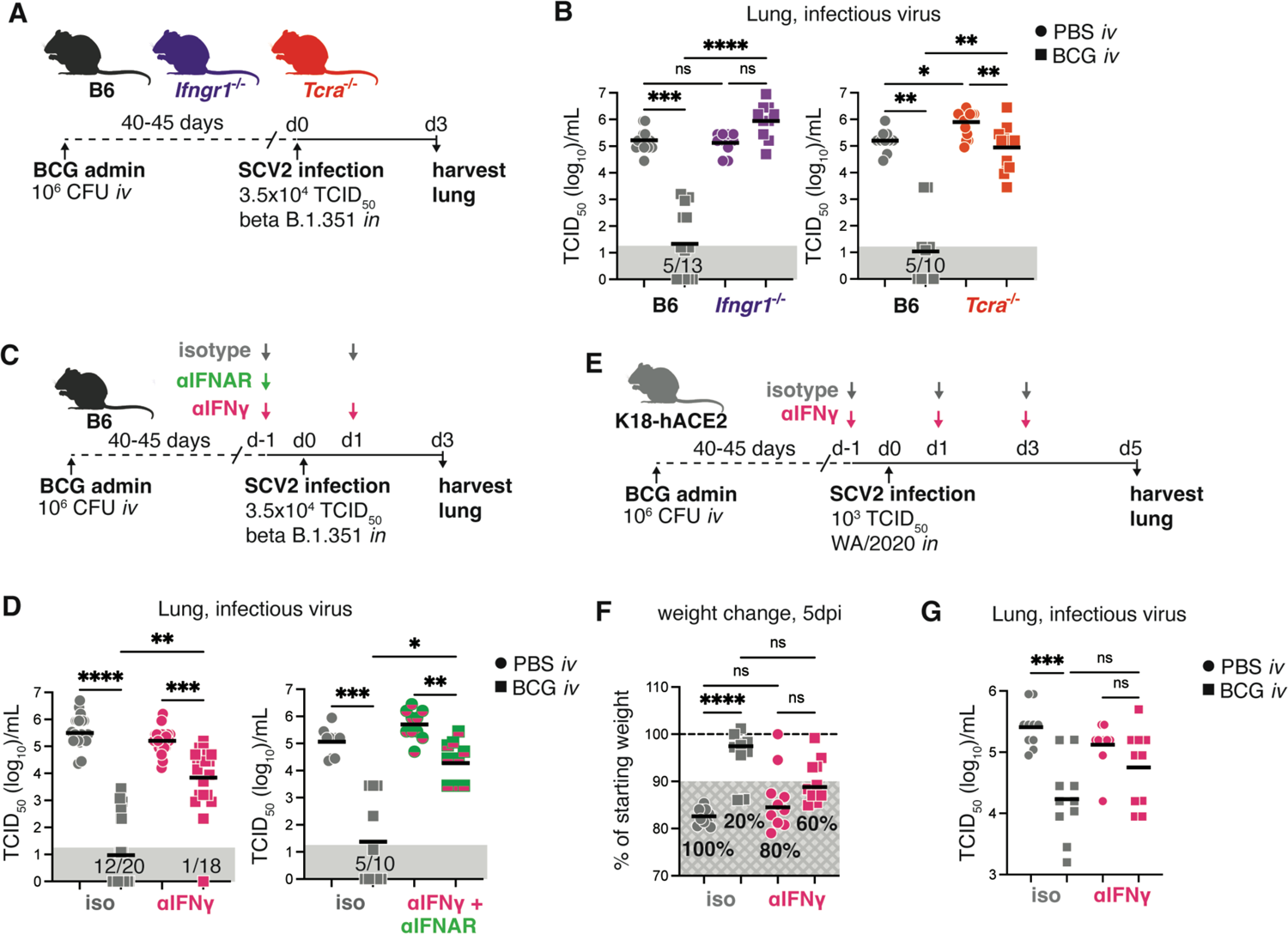
IFNγ is required for *iv* BCG-induced protection against early SCV2 infection. Mice of the indicated genotypes were inoculated with BCG or PBS *iv* 40-45 days prior to intranasal challenge with SCV2. Lungs were harvested 3-5 days after viral challenge. (**A**) Schematic of experimental protocol. (**B**) Viral titers in lung homogenate from B6, *Ifngr1*-/- or *Tcra*-/- mice as measured by TCID_50_ assay 3 days after SCV2 B.1.351 challenge. Data are pooled from 2 independent experiments with 4-5 mice/group. (**C**) Schematic of experimental protocol with anti-IFNγ and anti-IFNAR treatment. (**D**) Viral titers in lung homogenate as measured by TCID_50_ assay 3 days after SCV2 B.1.351 challenge. Data are pooled from 5 (anti-IFNγ only) or 2 (anti-IFNγ + anti-IFNAR) independent experiments with 4-5 mice/group. (**E**) Schematic of experimental protocol with anti-IFNγ treatment in the K18-hACE2 mouse model. (**F**) Percentage of starting weight at 5dpi. (**G**) Viral titers in lung homogenate as measured by TCID_50_ assay 5 days after SCV2 WA/2020 challenge. Data are pooled from 2 independent experiments with 4-5 mice/group. Statistical significance was assessed by Kruskal-Wallis with Dunn’s post-test. Not significant (ns) *p*>0.05; \**p*<0.05; **, *p*<0.01; \*\*\**p*<0.001; \*\*\*\**p*<0.0001. Gray boxes denote values below limit of detection.

We next tested whether IFNγ is required for *iv* BCG induced protection in transgenic K18-hACE2 mice which provide a model of SCV2-induced pathology and severe disease (**Fig 2E**) (*45*). *iv* BCG protected K18-hACE2 mice against weight loss and resulted in lower viral loads, but this protection was lost in animals treated with anti-IFNγ (**Fig 2F-G**). Together, these data from two different *in vivo* models demonstrate a key role for *iv* BCG-induced IFNγ in mediating protection against SCV2 infection. Furthermore, the observation that a short window of IFNγ neutralization is sufficient to significantly abrogate protection suggested that the cytokine mediates its effects at the time of initial SCV2 exposure.

### *iv* BCG-induced IFNγ promotes expression of viral restriction factors but is not required at the time of SCV2 challenge to limit virus induced inflammation

In addition to reducing viral loads, *iv* BCG limits SCV2 driven lethality and hyperinflammation (*33*). To examine whether IFNγ was also involved in *iv* BCG induced suppression of inflammation following SCV2, we performed a cytokine multiplex assay, flow cytometry and scRNAseq on lung samples from B6 mice 3 days after challenge with a SCV2 beta variant in the presence or absence of IFNγ neutralization (**Fig 3A**). While WT B6 mice do not develop lethal disease, they do respond with a characteristic SCV2 pro-inflammatory response consisting of heightened IFN-I, IFNλ, IL-6, GM-CSF and CCL2 levels (**Fig 3B-C**). These responses were absent or significantly lower in *iv* BCG inoculated animals irrespective of αIFNγ treatment (**Fig 3B**). Similar observations were apparent at the transcript level for *Il6*, *Csf2* (encoding GM-CSF), *Ccl2* and *Il18* across distinct cell lineages (**Fig 3C**), suggesting that dampening of SCV2-driven hyperinflammation by *iv* BCG occurs independently of IFNγ when this cytokine is neutralized at the time of viral challenge. This hypothesis was further supported by the reduction in virus driven tissue infiltrating inflammatory monocytes and their expression of the IFN inducible marker CD317 (Tetherin, encoded by the *Bst2* gene) in *iv* BCG inoculated mice with or without IFNγ neutralization (**Fig 3D**).

**Figure 3:**
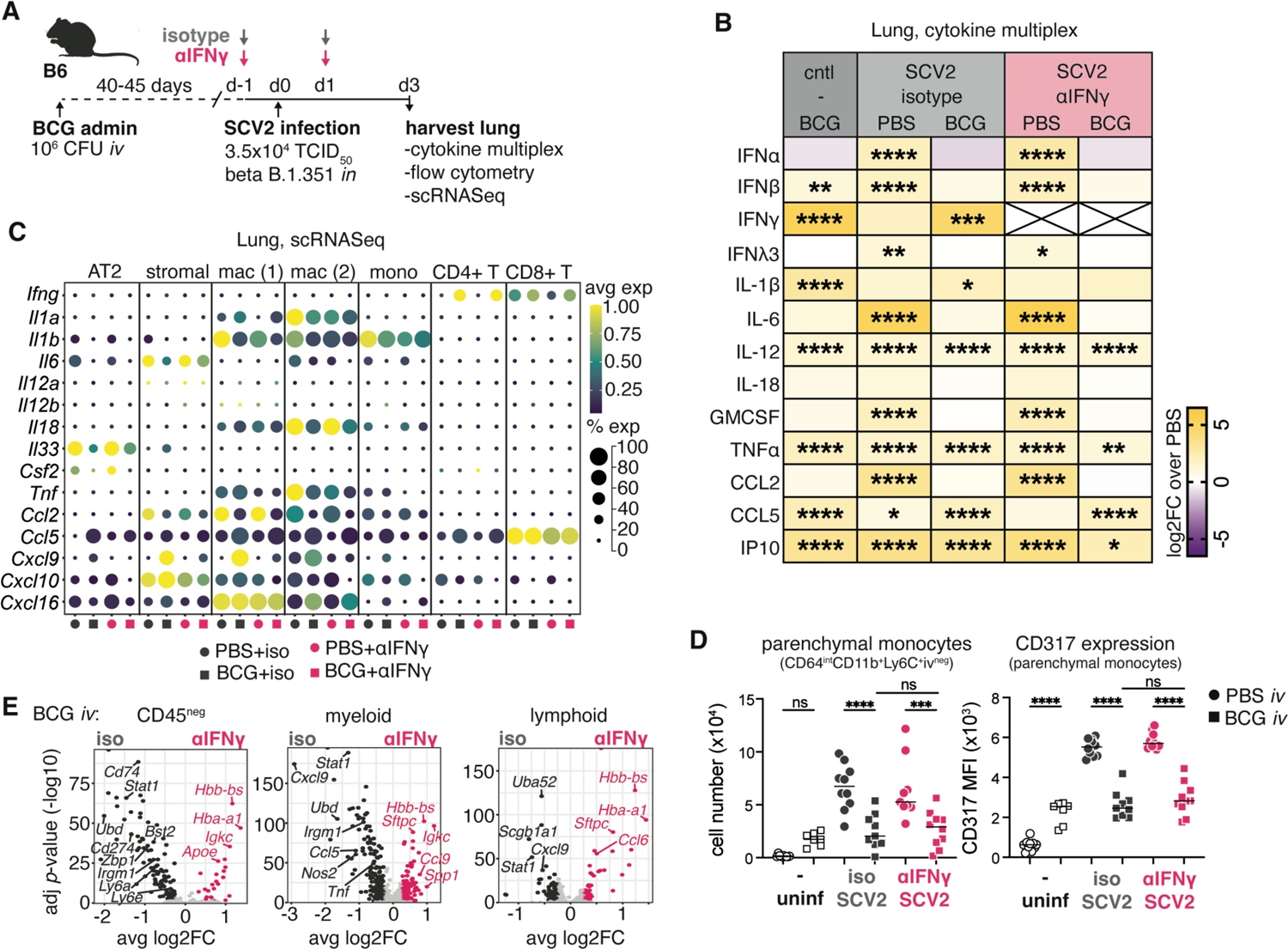
*iv* BCG-induced IFNγ present at the time of SCV2 challenge limits viral replication but not SCV2-induced inflammatory responses. B6 mice were inoculated with BCG or PBS *iv* 40-45 days prior to intranasal challenge with SCV2 B.1.351. Infected animals received an IFNγ neutralizing antibody or isotype control 1 day prior to and 1 day following SCV2 instillation. Lungs were harvested 3 days after viral challenge. (**A**) Schematic of experimental protocol. (**B**) Heat map display of log2 fold change (FC) of cytokine levels in lung homogenate relative to PBS uninfected controls. Cytokines were measured by multiplex assay and normalized to total protein content. Data are pooled from 3 independent experiments with 4-5 mice/group. Statistical significance was assessed by One-Way ANOVA with Tukey post-test. Not significant (ns) *p*>0.05; \**p*<0.05; **, *p*<0.01; \*\*\**p*<0.001; \*\*\*\**p*<0.0001. (**C**) Dot plot shows the relative expression and frequency of the indicated genes across selected Seurat clusters defined in the UMAP in Fig 1C. (**D**) The number of parenchymal monocytes (defined as live/CD45^+^/Ly6G^-^/CD64^int^/CD88^int^/CD26^-^/CD11b^+^/Ly6C^+^/iv^-^) and their expression of CD317 (MFI) as determined by flow cytometry. Data are pooled from 2 independent experiments with 4-5 mice/group. Statistical significance was assessed by One-Way ANOVA with Tukey post-test. Not significant (ns) *p*>0.05; \*\*\**p*<0.001; \*\*\*\**p*<0.0001. (**E**) Volcano plots show DEGs between isotype and anti-IFNγ treated mice inoculated *iv* with BCG across CD45^neg^, myeloid and lymphoid lineages that were manually annotated from the Seurat clustering showing in Fig 1C. Statistically significant DEGs are shown in dark gray or pink and were defined as FC>2.5 and *p*<0.05. Light grey points denote genes that did not reach statistical significance. Gene lists and their associated FC and *p*-values can be found in Supplementary Tables 5-7.

Interestingly, despite the 4-log fold increase in viral load observed after IFNγ blockade in BCG inoculated mice (**Fig 2D**), there were very few differences in the pulmonary cytokine milieu, other than the known IFNγ regulated cytokine, CXCL9 and TNFα (**Fig 3B-C**). *Tnf* transcript in monocytes was reduced upon IFNγ neutralization in BCG inoculated animals as were *Cxcl9* levels in macrophage and stromal populations (**Fig 3C**). Differential expression analysis of pooled CD45-negative, myeloid or lymphoid cells revealed an enrichment in ISGs (*Stat1*, *Ly6a*, *Irf1*, *Irgm1* and *Nos2*) in isotype treated BCG animals, confirming effective neutralization of IFNγ in mice treated with anti-IFNγ (**Fig 3E and Tables S5-7**). Notably, genes with known anti-SCV2 activity (*Bst2*, *Udb*, *Zbp1* and *Ly6e*) (*46–48*) were among the transcripts downregulated by IFNγ neutralization within the CD45-negative pool (**Fig 3E and Tables S5-7**). This downregulation of ISGs was not apparant in PBS controls treated with anti-IFNγ (**FigS3A and Tables S8-10**), consistent with the relatively low levels of IFNγ present during the early phase of SCV2 infection (**Fig 3B**). Overall, these results suggest that *iv* BCG induced IFNγ present at the time of viral challenge acts by directly controlling viral load rather than SCV2 driven inflammation.

### IFNγ receptor signaling is required in non-hematopoietic cells for *iv* BCG induced protection against SCV2

Given that SCV2 primarily infects pulmonary epithelial cells (EC) and that IFNγ neutralization impacts expression of anti-viral ISGs within the CD45-negative cellular compartment, we next wanted to determine whether IFNγ receptor signaling is specifically required in the non-hematopoietic compartment for *iv* BCG driven control of viral loads. To do this, *Ifngr1*-/- mice were lethally irradiated and reconstituted with bone marrow cells from WT B6 congenic donors, so that all radio-sensitive immune cells could signal through the IFNγ receptor while all radio-resistant cells, including the epithelial compartment, were *Ifngr1* deficient. To control for radiation induced stress and inflammation, we also generated WT B6 chimeras as comparators. Chimeras were injected *iv* with PBS or BCG and then challenged with SCV2 after 40-45 days (**Fig 4A**). As expected, lack of the IFNγ receptor on non-hematopoietic cells had no impact on SCV2 infection in the absence of BCG (**Fig 4B**). Viral loads were significantly lower in BCG inoculated control chimeras, but this protection was lost if the non-hematopoietic compartment was deficient in the IFNγ receptor indicating that BCG-induced IFNγ mediates control of viral loads through actions on non-hematopoietic cells (**Fig 4B**).

**Figure 4:**
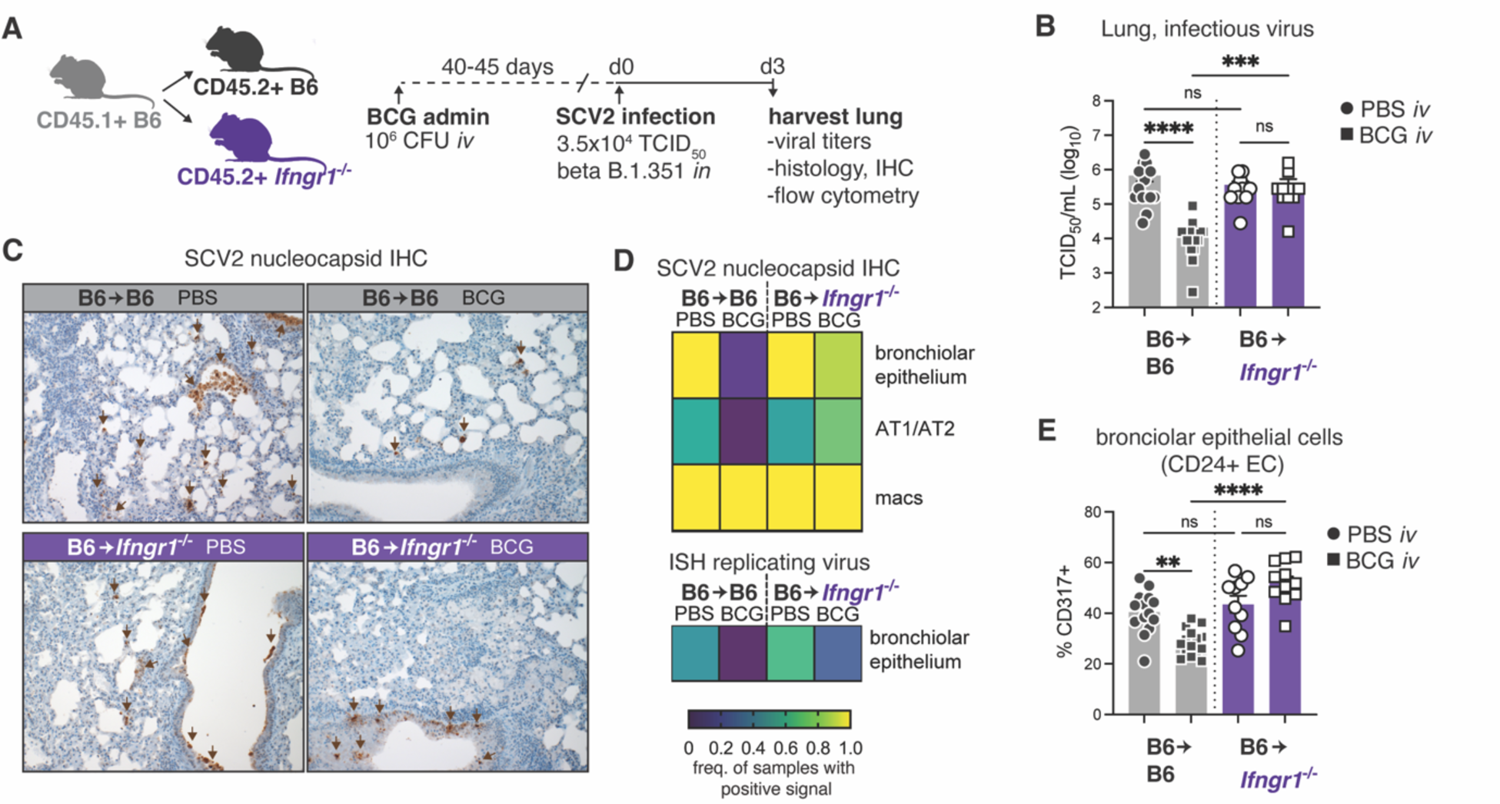
IFNγR1 signaling is required in the non-hematopoietic compartment for *iv* BCG induced protection against SCV2 infection. B6 congenic CD45.1+ mice were irradiated and reconstituted with either B6 CD45.2+ or *Ifngr1*-/- CD45.2+ bone marrow. Chimeras were inoculated with BCG or PBS *iv* 40-45 days prior to intranasal challenge with SCV2 B.1.351. Lungs were harvested 3 days after viral challenge. (**A**) Schematic of experimental protocol. (**B**) Viral titers in lung homogenate from B6 or *Ifngr1*-/- chimeras as measured by TCID_50_ assay. Data are pooled from 3 independent experiments with 3-5 mice/group. Statistical significance was assessed by Kruskal-Wallis with Dunn’s post-test. Not significant (ns) *p*>0.05; \*\*\**p*<0.001; \*\*\*\**p*<0.0001. (**C**) Representative lung histology images of SCV2 nucleocapsid immunohistochemical staining. (**D**) Heat map representation of SCV2 positivity across different cell types (IHC and ISH) as assessed by a study-blinded veterinary pathologist. Data are pooled from 2 independent experiments with 3-5 mice/group. (**E**) Expression of CD317 by CD24+ epithelial cell populations as determined by flow cytometry. Data are pooled from 3 independent experiments with 3-4 mice/group. Statistical significance was assessed by One-Way ANOVA with Tukey post-test. Not significant (ns) *p*>0.05; \**p*<0.05; **, *p*<0.01; \*\*\*\**p*<0.0001.

To more specifically characterize the impact of IFNγ on SCV2 infectivity of different cell types, we stained lung tissue sections for the SCV2 nucleocapsid and used *in situ* hybridization (ISH) targeting replicating viral RNA to identify actively infected cells (**Fig 4C and FigS4A-B**). We found that bronchiolar EC, pneumocytes and macrophages are the major cell types immunoreactive for the SCV2 nucleocapsid, with the bronchiolar epithelium the only lineage identified with active SCV2 infection by positive probe signal (ISH) at the experimental endpoint (**Fig 4D and FigS4B**). The immunoreactivity of macrophages is likely due to their role in their efferocytosis of debris of infected EC, rather than active infection (*49, 50*), explaining the high occurrence of positive signal observed in *iv* BCG inoculated control chimeras despite the low viral titer enumerated by TCID_50_ assay (**Fig 4C-D**). Importantly, lack of the IFNγ receptor on non-hematopoietic cells was associated with increased immunoreactivity for SCV2 nucleocapsid in bronchiolar EC and pneumocytes, as well as a higher frequency of rare ISH+ bronchiolar epithelial cells in *iv* BCG inoculated mice (**Fig 4C-D**). Consistent with these findings, flow cytometric analysis showed the IFN-inducible anti-viral response protein CD317 is more highly expressed by *Ifngr*-/- bronchial/bronchiolar EC (referred to as CD24+ EC) than *Ifngr*+/+ cells in *iv* BCG inoculated mice (**Fig 4E**), indicative of higher viral load. In these same animals, no differences were observed in CD317 expression by type-1 (AT1, CD326+ CD24-Pdpn+) or type-2 (AT2, CD326+ CD24-MHCII+) pneumocytes (**FigS4C**). Expression of CD274 (PDL1), which is strongly regulated by IFNγ, was significantly lower in all *Ifngr1*-/- epithelial cell types assessed from BCG inoculated animals confirming unresponsiveness of the epithelial compartment to bacteria-induced IFNγ (**FigS4D**). Together, these data support the conclusion that IFNγ produced following BCG injection controls SCV2 infectivity and/or replication within the epithelial compartment.

### IFNγ promotes expression of anti-viral proteins in pneumocytes and bronchiolar epithelial cells

To gain a deeper understanding of epithelial responses to BCG and SCV2, we assessed pulmonary EC by flow cytometry from control or *iv* BCG inoculated mice prior to or following SCV2 challenge (gating shown in **FigS5A**). We focused on expression of IFN inducible proteins with previously characterized roles in immune regulation (CD274, Ly6A/E) (*51, 52*) and anti-viral activity (CD317) (*46, 53, 54*). Interestingly, the expression pattern for each of the assessed proteins was distinct across different EC lineages (**Fig 5A and FigS5B-C**). BCG inoculation and SCV2 infection both drive strong up-regulation of IFN responsive proteins in AT1 and AT2 cells, with the anti-viral protein CD317 induced to similar levels by BCG and SCV2. CD24+ EC (bronchial/bronchiolar EC) also respond with upregulation of the assayed proteins following BCG and SCV2, although their response to SCV2 is much more pronounced than to BCG (**Fig 5A and FigS5B-C**). We identified a small subset of CD326+ CD24-Pdpn-MHCII-cells (referred to as CD24-EC) in our analysis although these cells show minimal responsiveness to BCG (**FigS5B-C**).

**Figure 5:**
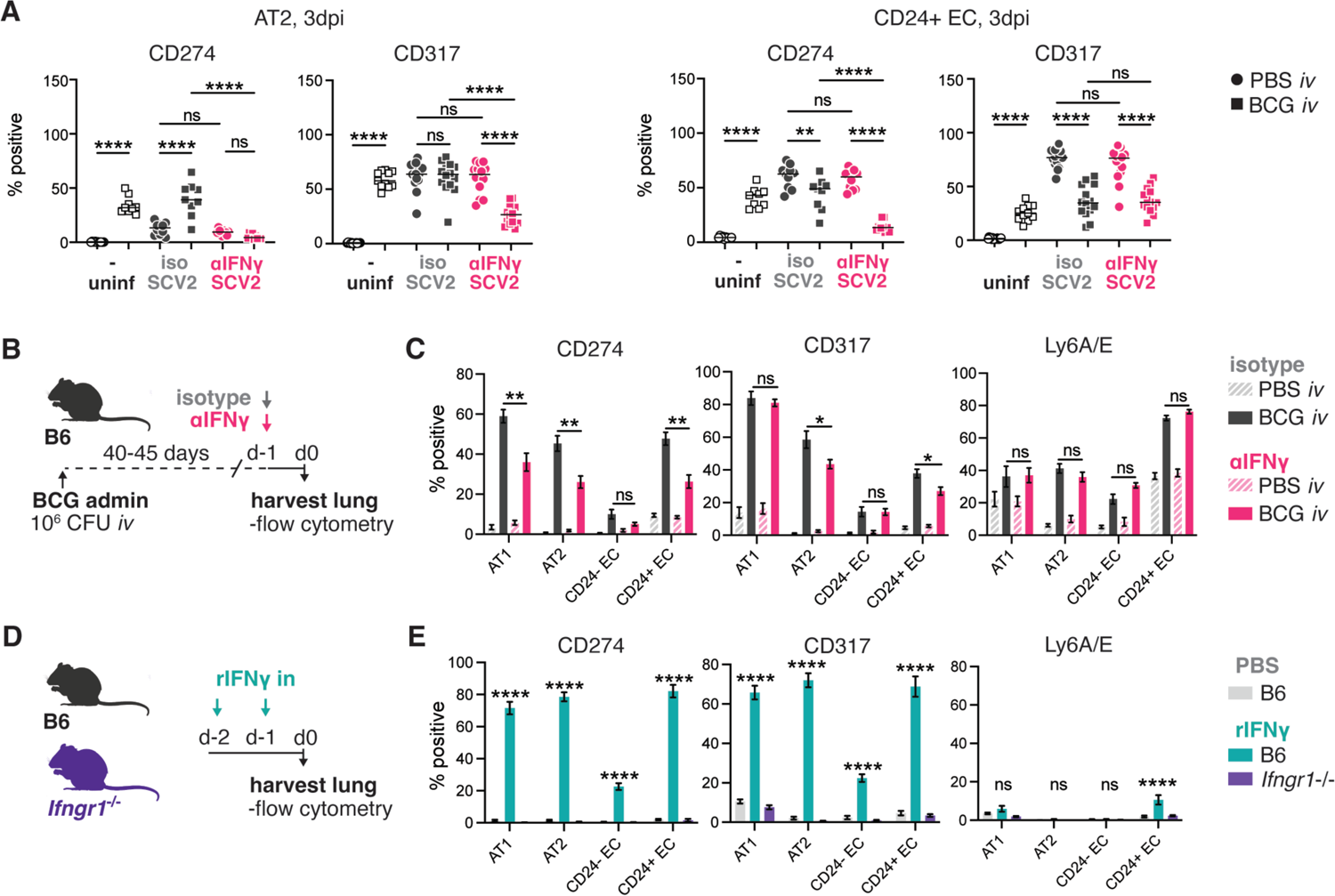
IFNγ induces expression of anti-viral markers in pneumocytes and CD24+ epithelial cells. (**A**) B6 mice were inoculated with BCG or PBS *iv* 40-45 days prior to intranasal challenge with SCV2 B.1.351. Infected animals received an IFNγ neutralizing antibody or isotype control 1 day prior to and 1 day following SCV2 instillation. Lungs were harvested 3 days after viral challenge and the indicated epithelial cell types assessed for CD274 and CD317 expression by flow cytometry. Data are pooled from 2-3 independent experiments with 4-5 mice/group. Statistical significance was assessed by One-Way ANOVA with Tukey post-test. Not significant (ns) *p*>0.05; \**p*<0.05; **, *p*<0.01; \*\*\*\**p*<0.0001. (**B-C**) B6 mice were inoculated with BCG or PBS *iv* 40-45 days prior to receiving an IFNγ neutralizing antibody or isotype control. Lungs were harvested 1 day after anti-IFNγ treatment. (**B**) Schematic of experimental outline. (**C**) Expression of CD274, CD317 and Ly6A/E across different epithelial cell types. Data are pooled from 2 independent experiments with 3-4 mice/group. Statistical significance was assessed by unpaired t-test between BCG isotype and BCG anti-IFNγ for each cell type. Not significant (ns) *p*>0.05; \**p*<0.05; **, *p*<0.01; \*\*\**p*<0.001. (**D-E**) B6 or *Ifngr1*-/- mice were treated with PBS or rIFNγ intranasally on 2 consecutive days. Lungs were harvested 1 day after the last treatment. (**D**) Schematic of experimental protocol. (**E**) Expression of CD274, CD317 and Ly6A/E across different epithelial cell types. Data are pooled from 2 independent experiments with 4-5 mice/group. Statistical significance was assessed by unpaired t-test between PBS and rIFNγ treated B6 mice for each cell type. Not significant (ns) *p*>0.05; \*\*\*\**p*<0.0001.

Given the ability of BCG to induce expression of proteins involved in anti-SCV2 activity, we hypothesized that mycobacterial-induced IFNγ may be inducing an “anti-viral” state in pulmonary EC prior to SCV2 challenge thus limiting the infectivity of the virus. To examine this possibility, we inoculated animals with BCG and then neutralized IFNγ after 40-45 days before assessing the pulmonary EC 1 day later (ie. at the time we would usually challenge with SCV2) (**Fig 5B**). Again, we observed distinct expression patterns for CD274, CD317 and Ly6A/E across the different epithelial subsets. CD274 expression was the most strongly impacted by IFNγ neutralization, which was not unexpected given its well documented regulation by IFNγ (*55*). Ly6A/E expression was not impacted by anti-IFNγ treatment, but CD317 was significantly reduced, albeit modestly, in AT2 and CD24+ EC, the two major cell types that are targeted for infection by SCV2 (**Fig 5C**). We next performed an experiment administering recombinant (r)IFNγ intranasally to naïve WT mice or animals that report expression of the IFN-inducible protein Irgm1 (M1Red) and then assessed EC responses by flow cytometry (**Fig 5D and FigS5D**). These data closely match the observed epithelial response following *iv* BCG, with pneumocytes and CD24+ EC strongly upregulating CD274, CD317 and Irgm1 after rIFNγ treatment and only showing low level expression of Ly6A/E **(Fig 5E and FigS5E-F)**. As observed with BCG, CD24-EC had a lower response to IFNγ compared to the other EC subsets **(Fig 5E)**, except for Irgm which was similarly induced across all cell types following rIFNγ treatment (**FigS5E-F)**. Together, our data demonstrate that BCG-driven IFNγ induces expression of IFN-regulated proteins by pulmonary EC, including the viral restriction factor CD317.

### Intranasal administration of recombinant IFNγ confers strong protection against SCV2 in two mouse models

To test whether IFNγ is sufficient to confer protection against SCV2 in the absence of BCG inoculation, we administered the recombinant cytokine intranasally to WT or *Ifngr1*-/- animals on the two days preceding and the day following SCV2 challenge (**Fig 6A**). At 3 days post infection, viral titers in the lung tissue were significantly lower in WT B6 animals that received intranasal rIFNγ, with more than 50% of treated animals having no detectable infectious particles at this timepoint (**Fig 6B**). Importantly, no protection was observed in rIFNγ-treated *Ifngr1*-/- mice confirming that rIFNγ exerted anti-viral activities through its receptor (**Fig 6B**). Immunohistochemical sanalysis of the SCV2 nucleocapsid in lung sections showed a very similar picture, with reduced viral immunoreactivty in lungs from rIFNγ-treated WT B6 mice across all cell types assessed (**Fig 6C-D**). Notably, intranasal rIFNγ also significantly protected against SCV2 induced pulmonary inflammation and pneumonia in this mild infection model (**Fig 6C, E**), indicating that it may also be protective against severe disease which is characterized by marked tissue chages and lung damage (*45*).

**Figure 6:**
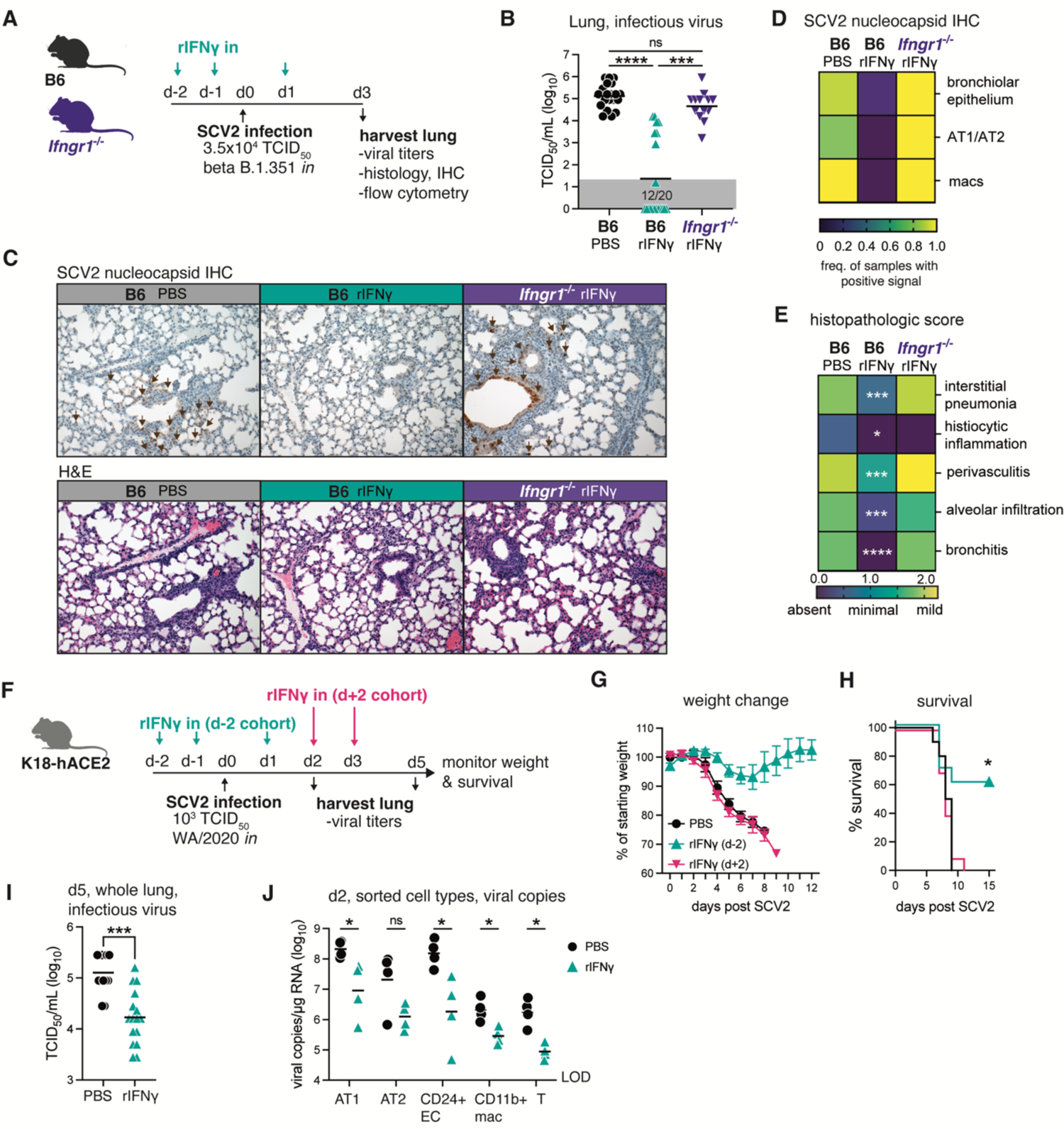
Intranasal administration of recombinant IFNγ prior to viral challenge confers strong protection against SCV2. (**A-E**) B6 or *Ifngr1*-/- mice were infected with SCV2 B.1.351 and lungs harvested for analysis 3dpi. Animals were treated with PBS or rIFNγ intranasally on days −2, −1 and 1 relative to viral challenge. (**A**) Schematic of experimental protocol. (**B**) Viral titers in lung homogenate as measured by TCID50 assay. Data are pooled from 4 independent experiments with 4-5 mice/group. Statistical significance was assessed by Kruskal-Wallis with Dunn’s post-test. Not significant (ns) *p*>0.05; \*\*\**p*<0.001; \*\*\*\**p*<0.0001. Gray boxes denote values below limit of detection. (**C**) Representative lung histology images stained for SCV2 nucleocapsid (upper panel) or H&E (lower panel). (**D-E**) Heat map representation of SCV2 positivity across different cell types (**D**) or histopathologic score (**E**) as assessed by a study-blinded veterinary pathologist. (**F-J**) K18-hACE2 mice were infected with SCV2 WA/2020. Animals received intranasal PBS or rIFNγ at the indicated time points. (**F**) Schematic of experimental protocol. (**G**) Percent of starting weight and (**H**) survival over time following viral challenge. Data are pooled from 2 independent experiments with 5 mice/group. Statistical significance was determined using a Mantel-Cox test. \**p*<0.05. (**I**) Viral titer in lung homogenate 5 dpi as measured by TCID50 assay. Data are pooled from 2 independent experiments with 7-8 mice/group. Statistical significance was determined by Mann-Whitney test. \*\*\**p*<0.001. (**J**) Single cell suspensions were prepared from lungs at 2dpi. Cells were pooled into 2 replicates/experimental treatment group each containing cells from 2-3 mice. The indicated cell types were sorted as per the gating strategy in FigS6. RNA was directly extracted, and viral copies measured by PCR. Data are pooled from 2 independent experiments. Statistical significance was determined by Mann Whitney test for each cell type. Not significant (ns) *p*>0.05; \**p*<0.05. LOD = level of detection.

To address the latter hypothesis, we performed an additional experiment in which we treated highly susceptible K18-hACE2 mice intranasally with rIFNγ prior to challenging with SCV2 (rIFNγ d-2) and then assessed survival, body weight and viral titers (**Fig 6F**). Based on our analysis of EC (**Fig 5**), we hypothesized that IFNγ predominantly exerts its protective effects through induction of anti-viral programs in cells targeted for SCV2 infection prior to viral exposure. To examine this possibility, we also included a group of mice in the survival study that started rIFNγ treatment at the peak of viral replication, 2 days after SCV2 challenge (rIFNγ d+2) (**Fig 6F**). Animals that received rIFNγ from d+2 and the PBS control group rapidly lost weight following SCV2 exposure (**Fig 6G**), with all mice succumbing to infection by 11 days post challenge (**Fig 6H**). In contrast, rIFNγ treatment starting prior to SCV2 protected against weight loss in the majority of animals and significantly improved survival (**Fig 6G-H**), suggesting that IFNγ-induced responses are required at-or-near the time of viral exposure to be prevent the establishment of SCV2 infection and protect against disease. Viral titers assayed in whole lung homogenate at 5dpi showed lower SCV2 burdens with rIFNγ treatment, consistent with our findings in the B6 model (**Fig 6I**). Finally, we assessed viral loads in individual cell types sorted from SCV2-infected K18-hACE2 mice 2 days after challenge to determine whether rIFNγ preferentially affected anti-viral activity in a particular cell type (sorting strategy shown in **FigS6**). As expected, EC subsets (AT1, AT2 and CD24+ EC) had the highest number of viral copies per µg of RNA, suggestive of active infection (**Fig 6J**). SCV2 copies were also detected at low levels in sorted macrophages and T cells, which likely resulted from “sticky” viral material released from surrounding dead/dying infected cells. In all cell types assessed, rIFNγ treatment reduced the number of viral copies recovered; however, this did not reach statistical significance in the AT2 group. Overall, these data demonstrate that pre-existing IFNγ responses limit SCV2 infection and/or replication across several epithelial cell lineages thus protecting the host from virus-induced tissue damage and immunopathology.

## Discussion

Rapid and robust type-1 or type-3 IFN responses are crucial for effective control of viruses. One feature that makes SCV2 such a successful pathogen is its ability to suppress host IFN responses and MHCI expression, much more so than other common respiratory viruses such as influenza A (*56–59*). It is this immunosuppressive property that potentially makes SCV2 more amenable to restriction by pre-established IFN responses such as those driven by concurrent or recent pulmonary infections. We have previously reported that pulmonary infection with BCG achieved through *iv* inoculation provides striking protection against SCV2 driven pathology and lethality, whereas subcutaneous administration does not (*33*). In the present study we investigated the mechanism underlying this BCG-induced anti-viral resistance and formally demonstrate that IFNγ (type-2 IFN) signaling, specifically in non-hematopoietic cells, is essential for the observed protection against SCV2. We further demonstrate that intranasal treatment with the recombinant cytokine restricts SCV2 infection/replication and can protect the host from lethal disease.

Our findings reveal that pre-existing IFNγ can substitute for early type-1 or type-3 responses in controlling SCV2 infection. However, IFNγ does not appear to play a role during the natural course of SCV2 infection since as shown here animals deficient in IFNγ receptor signaling do not display impaired viral control in the absence of BCG. This conclusion is in agreement with previous findings in both mice (*25, 60*) and rhesus macaques (*61*). Furthermore, while deficiencies in IFNγ signaling have been characterized in humans and linked with increased susceptibility to some viral pathogens (*62*), there is little evidence associating such mutations with increased risk of COVID-19 despite intensive investigation in this area. Rather, inborn errors in the IFN-I pathway are strongly associated with severe COVID-19 (*11, 12*) as well as other viral infections (*63*), whereas IFNγ signaling deficiencies are often linked with susceptibility to mycobacterial disease (*64, 65*). Redundant functions by type-1 and type-3 IFNs that are induced earlier during the course of viral infection are a potential explanation for lack of IFNγ requirement in these settings. Indeed, combined IFN receptor deficiencies (eg. *Ifngr1* and *Ifnar1*) render mice highly susceptible to SCV2 (*25, 66*). Furthermore, in the case of influenza, IFNγ responses have been linked to protection from secondary viral challenge as primary infection induced tissue-resident memory T cells are able to provide IFNγ rapidly upon re-exposure (*18*).

The above discussion highlights the importance of prompt IFNγ induction relative to the timing of viral exposure for beneficial outcomes. We directly tested this requirement in our study by comparing survival between animals treated with rIFNγ 2 days prior to versus 2 days following SCV2 infection and found only pre-existing IFNγ responses could protect mice. Similar conclusions have been made in other animal studies assessing prophylactic versus therapeutic administration of with recombinant type-1 or type-3 IFNs (*25, 30*) or a type-1 IFN-inducing RIG-I agonist (*31*). Interestingly, therapeutic treatment with IFNλ starting less than 24 hours following SCV2 infection provided some protection in susceptible aged BALB/c or *Ifnar*-/- mice (*25, 67*) that was significantly enhanced if the IFNλ treatment was combined with rIFNγ (*25*). This finding suggests that combination treatments that include IFNγ could be effective in a therapeutic setting. In this regard, Beer *et al*. speculate that in their aged mouse model, IFNλ provides anti-viral signals whereas IFNγ restores an age-related delay in immune cell recruitment, which may explain why there was limited efficacy when each cytokine was administered individually. Our data suggest that IFNγ also contributes directly to viral load control, particularly if it is delivered intranasally (as in the current study) versus subcutaneously (*25*).

The presence of an IFN response within the pulmonary compartment at the time of SCV2 exposure is most likely to occur in the context of an ongoing or recent viral or bacterial infection. We show here that pulmonary BCG infection achieved through *iv* administration induces a strong IFNγ signature in both myeloid and epithelial cells and that blockade of this response reverses protection against SCV2. Our scRNAseq and flow cytometry data demonstrate that *iv* BCG has a pronounced impact on the transcriptional landscape of pulmonary immune cells, with the replacement of resident alveolar macrophages by IFNγ primed monocyte-derived cells and the recruitment of Th1 cells being the most prominent changes. IFNγ transcript and protein were highly expressed by Th1 cells and *iv* BCG mice that lacked T cells were unable to control SCV2 infection as effectively as their WT counterparts. T cell deficiency had no significant impact on viral titers at 3dpi in the absence of BCG, suggesting that the effect observed was due to BCG-induced T cell derived IFNγ. Another interesting note from this experiment was that despite carrying a substantial bacterial load, *Tcra*-/- and *Ifngr1*-/- mice were still susceptible to SCV2 suggesting that the mere presence of an ongoing BCG infection is insufficient to limit viral infection. This observation could explain the discrepancy in the lack of *iv* BCG induced resistance in SCV2-infected rodents reported by Kaufmann *et al*. (*68*) and the protection observed by us and other investigators (*33–35*) since failure to induce a sufficient IFNγ response following *iv* BCG administration due to potential differences in bacterial strain, preparation or dosing would not result in protection.

Importantly, in our murine models IFNγ mediated protection against SCV2 occurs within the non-hematopoietic compartment. Given that pulmonary epithelial cells (ECs) are the primary target for SCV2 infection (*3–5*) and that IFNγR-deficient ECs in the presence of a IFNγR-sufficient immune compartment still have increased viral loads, we propose that bacteria-induced IFNγ likely mediates its anti-viral effects directly within EC. However, without cell-type specific deletion of the *Ifngr1*, we cannot rule out contribution of other radioresistant cells. EC and other cells targeted by viruses combat infection through IFN regulated induction of host-derived viral restriction factors that are among a broader group of IFN-stimulated genes (ISGs) (*6*). While thousands of ISGs have been identified (*16*), the precise mechanisms of anti-viral activity have only been described for a few dozen, which are typically regulated by type-1 and type-3 IFNs (*6*). One such protein, CD317 (Tetherin, encoded by the *Bst2* gene), is expressed on the cell surface and interferes with the release of viral particles, including SCV2, from infected cells (*46, 53, 54*). CD317 is strongly induced by type-1 IFN but is also reported to be up-regulated by IFNγ depending on the cell type assessed (*16, 54, 69*). We show here that IFNγ promotes CD317 expression by pulmonary ECs and that the level induced by BCG infection is equivalent to that induced on AT2 cells by SCV2 exposure. These data align with previous observations that pulmonary EC are highly responsive to rIFNγ treatment and influenza induced IFNγ (as measured by Irgm1-DSRed expression) (*70*). Together the above findings suggest that IFNγ can directly stimulate and induce expression of ISGs in pulmonary epithelial cells that may be involved in restricting early viral replication and identify CD317 as a candidate for further study into IFNγ conferred anti-viral activity.

The major role of IFNγ in mediating host protection against SCV2 appears to be through controlling viral load, which leads to lower levels of virus induced pathology and mortality. This finding is clear in the rIFNγ model where viral loads were lower, lung pathology was reduced, and animals had improved survival. Interestingly, our scRNAseq, flow cytometry and cytokine multiplex data from *iv* BCG animals infected with SCV2 showed that despite the higher viral load following IFNγ neutralization, markers of inflammation including IL-6 and CCL2 production were still reduced, along with less accumulation of inflammatory monocytes in the lung tissue compared to PBS controls. This suggests that other factors may be playing a role in addition to IFNγ in dampening virus-induced inflammation in the lungs of mice concurrently infected with BCG. An alternative possibility is that the short window of IFNγ neutralization employed in these studies was enough to impact viral loads but that residual IFNγ imprinting of the epithelium and other cell types was sufficient to protect against inflammatory cytokine production and immune cell recruitment.

Overall, our findings indicate that bacterial infections that specifically induce IFNγ responses within the lung may restrict SCV2 infection. Interactions with other bacterial pathogens such as *Streptococcus pneumoniae* (Spn) that drive predominant Th17 responses may not be protective, with one mouse study showing that Spn colonized animals are more susceptible to SCV2 infection (*71*). Meanwhile, aerosol infection with *Mycobacterium tuberculosis* (Mtb) does protect mice against SCV2 (*36, 37*) independent of type-1 IFN signaling (Paul Baker and Katrin Mayer-Barber, personal communication). Like BCG, Mtb invokes a strong IFNγ response within the lung that potentially plays a part in the observed anti-viral effect. The fact that escalating Mtb dose increasingly restricts SCV2 (Paul Baker and Katrin Mayer-Barber, personal communication) further supports a possible role for IFNγ in the murine models of Mtb mediated SCV2 restriction. Despite also driving strong IFNγ responses in humans, Mtb infected individuals are not protected from SCV2 and if anything, appear to show increased COVID-19 disease (*72, 73*). The reasons behind the discrepancy between the mouse and human data is likely multifaceted and related to a number of epidemiological and socioeconomic factors in addition to biological ones. It is clear however that IFNγ does possess anti-viral properties in humans, with rIFNγ treatment of a human pulmonary epithelial cell line effective at limiting SCV2 infection *in vitro* (*59, 74, 75*) and subcutaneous rIFNγ treatment of a small group of moderately ill COVID-19 patients associated with reduced time to hospital discharge (*29*). The work presented here raises the possibility that prophylactic intranasal rIFNγ administration could protect exposed individuals against SCV2 infection or perhaps enhance the efficacy of other IFN treatments (eg. pegylated IFNλ).

## Methods

### Study design

The aim of this study was the elucidate the mechanism/s by which concurrent mycobacterial infection protects against SARS-CoV-2. Two mouse models of SCV2 infection were employed in this study: 1) commercially available K18-hACE2 transgenic mice that exhibit high suseptibility to SCV2 and severe pathology (*45*) and 2) non-transgenic mice, either wildtype or gene knockouts, that display a mild disease phenotype when infected with SCV2 B.1.351 (*38–42*). Animals were randomly assigned to groups of 3-8 mice for each experiment. Data from all experiments were pooled prior to analysis to identify reproducible and statistically significant differences between experimental groups. The number of mice per group, the number of experimental replicates and the statistical tests employed are reported in the figure legends. All data points are biological replicates. No animals were excluded from analysis except for technical failure of intranasal inoculation. Endpoint criteria for survival studies were pre-determined in line with animal welfare recommendations set by the NIAID Animal Care and Use Committee.

### Mice

C57BL/6J (JAX664), B6(Cg)-Ifnar1tm1.2Ees/J (JAX28288) and B6.Cg-Tg(K18-ACE2)2Prlmn/J hemizygous (JAX34860) mice were purchased from The Jackson Laboratory (Bar Harbor, ME); B6.PL-Thy1a/CyJ (JAX406), B6.SJL-Ptprca Pepcb/BoyJ (JAX2014) and B6.129S7-Ifngr1tm1Agt/J (JAX3288) mice were acquired from the NIAID Contract Facility at Taconic Farms; M1Red mice (*70*) were bred onsite at NIAID. Mice were housed under specific pathogen–free conditions with *ad libitum* access to food and water and were randomly assigned to sex- and age-matched experimental groups. All animal studies were conducted in AALAC–accredited Biosafety Level 2 and 3 facilities at the NIAID, National Institutes of Health (NIH) in accordance with protocols approved by the NIAID Animal Care and Use Committee.

To generate bone marrow chimeras, CD45.2+ B6 and *Ifngr1*^-/-^ mice received to two doses of 500cGy gamma-radiation, with a three-hour rest period between exposures. The following day, 10^7^ bone marrow cells from CD45.1+ B6.SJL donors were administered by intravenous injection. Animals were maintained on antibiotic drinking water for 3 weeks and rested for a further 5 weeks before the commencement of experiments. Flow cytometry was performed on peripheral blood cells to confirm successful reconstitution.

### Virology

SARS-CoV-2 strains USA-WA1/2020 (BEI Resources) and RSA B.1.351 N501Y (BEI Resources) were propagated in Vero-TMPRSS2 cells (kindly provided by Dr. Jonathan Yewdell, NIAID). Vero-TMPRSS2 cells were maintained in DMEM medium supplemented with glutamax, 10% FCS and 250µg/ml Hygromycin B gold (InvivoGen). Virus stock production was performed under BSL-3 conditions using DMEM medium supplemented with glutamax and 2% FCS. At 48h post inoculation, culture supernatant and cells were collected, clarified by centrifugation for 10 min at 4°C. Supernatant was collected, aliquoted and frozen at −80°C. Viral titers were determined by TCID_50_ assay in Vero E6 cells (ATCC CRL-1586) using the Reed and Muench calculation method. Full genome sequencing was performed at the NIAID Genomic Core (Hamilton, MT).

### BCG

BCG Pasteur from the Trudeau Collection was originally obtained from Dr. Sheldon Morris (Food and Drug Administration, MD) and maintained as laboratory stock by serial passage. BCG was propagated in 7H9 broth supplemented with 10% oleic acid-albumin-dextrose-catalase (OADC) enrichment media (BD Biosciences) until mid-log phase. Bacteria were harvested, washed thrice and frozen down in aliquots until use. Colony forming units were enumerated by culturing on 7H11 agar for 3 weeks at 37°C.

### Infections and treatments

BCG was prepared in PBS containing 0.05% Tween-80. A dose of 10^6^ CFU/mouse in 100µL was delivered by intravenous injection. Control animals received the same volume of PBS with 0.05% Tween80.

Recombinant murine IFNγ (R&D Systems) was reconstituted in PBS containing 0.01% normal mouse serum, aliquoted and frozen at −80°C until use. Just prior to administration, aliquots were thawed and diluted in PBS. Mice were anesthetized by isoflurane inhalation and 1µg rIFNγ in a volume of 35µL was administered by intranasal instillation on days −2, −1 and 1 as indicated in the text and figures. Control animals received 35µL PBS intranasally.

Anti-IFNγ (XMG1.2) or rat IgG1 isotype control (HPRN) was administered by intraperitoneal injection on days −1 (750µg) and 1 (250µg) as indicated in the text and figures. In some experiments, K18-hACE2 mice received an additional 250µg dose on day 3. Anti-IFNAR (MAR1-5A3) or mouse IgG1 isotype control (MOPC-21) was administered by intraperitoneal injection on day −1 (2mg) as indicated in the related figure. Antibodies were stored at 4°C until use and diluted in PBS just prior to administration. All antibodies were from BioXCell.

SCV2 infections were performed under BSL3 containment. Animals were anesthetized by isoflurane inhalation and a dose of 10^3^ TCID_50_/mouse SCV2 WA/2020 or 3.5×10^4^ TCID_50_/mouse SCV2 B.1.351 was administered by intranasal instillation. Following infection, mice were monitored daily for weight change and clinical signs of disease by a study-blinded observer.

### Determination of viral copies by quantitative PCR

Lung and brain were homogenized in Trizol and RNA was extracted using the Direct-zol RNA Miniprep kit following the manufacturer’s instructions. E gene gRNA was detected using the QuantiNova Probe RT-PCR Kit and protocol and primers (forward primer: 5’-ACAGGTACGTTAATAGTTAATAGCGT-3’, reverse primer:5’-ATATTGCAGCAGTACGCACACA-3’) and probe (5′-FAM-ACACTAGCCATCCTTACTGCGCTTCG-3IABkFQ-3′) as previously described (*76*). The standard curve for each PCR run was generated using the inactivated SARS-CoV-2 RNA obtained from BEI (NR-52347) to calculate the viral copy number in the samples. Identical lung and brain portions were utilized for all experiments to generate comparable results.

### Determination of viral titers by TCID_50_ assay

Viral titers from lung and brain homogenate were determined by plating in triplicate on Vero E6 cells (kindly provided by Dr. Sonja Best, NIAID) using 10-fold serial dilutions. Plates were stained with crystal violet after 96 hours to assess cytopathic effect (CPE). Viral titers were determined using the Reed-Muench method (*77*).

### Determination of colony forming units

Bacterial burdens were enumerated by plating serially diluted lung homogenate on 7H11 agar (Sigma Aldrich) supplemented with 0.5% glycerol (Sigma Aldrich) and 10% OADC. BCG colonies were counted after a 3-week incubation at 37°C.

### Preparation of single cell suspensions from lungs

Lung lobes were diced into small pieces and incubated in RPMI containing 0.33mg/mL Liberase TL and 0.1mg/mL DNase I (both from Sigma Aldrich) at 37°C for 45 minutes under agitation (200rpm). Enzymatic activity was stopped by adding FCS. Digested lung was filtered through a 70µm cell strainer and washed with RPMI. Red blood cells were lysed with the addition of ammonium-chloride-potassium buffer (Gibco) for 3 minutes at room temperature. Cells were then washed with RPMI supplemented with 10% FCS. Live cell numbers were enumerated using AOPI staining on a Cellometer Auto 2000 Cell Counter (Nexcelom).

### Flow cytometry

To label cells within the pulmonary vasculature for flow cytometric analysis, 2µg anti-CD45 (30-F11; Invitrogen) was administered by intravenous injection 3 minutes prior to euthanasia.

Single-cell suspensions prepared from lungs were washed twice with PBS prior to incubating with Zombie UV™ Fixable Viability Dye and TruStain FcX™ (clone 93; both from BioLegend) for 15 minutes at room temperature. Cocktails of fluorescently conjugated antibodies diluted in PBS and 10% Brilliant Stain Buffer (BD) were then added directly to cells and incubated for a further 20 minutes at room temperature. Anti-CD11b (M1/70) and anti-CD26 (H194-112) were from BD OptiBuild. Anti-CD4 (GK1.5), anti-CD19 (GL3), anti-CD24 (M1/69), anti-CD44 (IM7), anti-CD45 (30-F11), anti-Siglec F (E50-2440), anti-TCR-beta chain (H57-597) and anti-TCR-gamma-delta (GL3) were from BD Horizon. Anti-CD8-alpha (53-6.7) and anti-CD274 (PDL1, clone MIH5) were from Invitrogen. Anti-CD11c (N418), anti-CD19 (6D5), anti-CD31 (390), anti-CD44 (IM7), anti-CD49f (GoH3), anti-CD64 (X54-5/7.1), anti-CD69 (H1.2F3), anti-CD86 (GL-1), anti-CD88 (20/70), anti-CD90.2 (30-H12), anti-CD104 (346-11A), anti-CD140a (APA5), anti-CD279 (PD1, clone 29F.1A12), anti-CD317 (BST2, clone 927), anti-CD326 (Epcam, clone G8.8), anti-IA/IE (MHCII, clone M5/114), anti-Ly6A/E (Sca-1, clone D7), anti-Ly6C (HK1.4), anti-Ly6G (1A8), anti-NK1.1 (PK136), anti-podoplanin (8.1.1), anti-TCR-beta chain (H57-597), anti-TCR-gamma-delta (GL3) and anti-XCR1 (ZET) were from BioLegend.

Cells were incubated in eBioscience™ Transcription Factor Fixation and Permeabilization solution (Invitrogen) for 2-18 hours at 4°C and stained with cocktails of fluorescently-labeled antibodies against intracellular antigens diluted in Permeabilization Buffer (Invitrogen) for 30 minutes at 4°C. Anti-Granzyme B (GB11) and anti-Ki67 (B56) were from BD Pharmingen. Anti-FoxP3 (FJK-16s), anti-NOS2 (CXNFT) and anti-Tbet (4B10) were from Invitrogen.

Compensation was set in each experiment using UltraComp eBeads^™^ (Invitrogen) and dead cells and doublets were excluded from analysis. All samples were collected on a FACSymphony A5 SORP^™^ flow cytometer (BD) and analyzed using FlowJo software (version 10, BD) (*78*).

### Cell sorting

Single-cell suspensions prepared from lungs were pooled from 2-3 mice into 2 replicates per experimental group. Cells were incubated in TruStain FcX™ (clone 93; BioLegend) diluted in PBS for 10 minutes at 4°C and subsequently stained with fluorescently conjugated antibodies diluted in PBS and 10% FCS for a further 30 minutes at 4°C. Anti-CD11b (M1/70), anti-CD24 (M1-69), anti-CD31 (390), anti-CD88 (20/70), anti-CD104 (346-11A), anti-CD326 (Epcam, clone G8.8), anti-IA/IE (MHCII, clone M5/114), anti-Ly6G (1A8), anti-podoplanin (8.1.1) and anti-TCR-beta chain (H57-597) were from BioLegend. Anti-CD45 (30-F11) and anti-F4/80 (BM8) were from Invitrogen. Following staining, cells were washed twice with PBS containing 5% FCS and stored on ice until sorting. Propidium iodide (Thermofisher) was added to samples just prior to sorting to exclude dead cells. Populations of interest were sorted under BSL3 containment on a FACSAria^™^ III cell sorter (BD) fitted with a 100µm nozzle into PBS containing 20% FCS. The gating strategy is shown in **FigS6**.

### Multiplex Cytokine Array and ELISA

Cytokines were assessed in lung homogenate using a ProcartaPlex Luminex kit (ThermoFisher) according to the manufacturers’ instructions and measured using a MagPix Instrument (R&D Systems). Interferon-lambda-3 was measured by Duoset ELISA (R&D Systems). Total protein was determined by BCA Assay (ThermoFisher). Cytokine levels were standardized to total protein content.

### Single-cell RNA sequencing

Single cell suspensions were prepared from lungs as described above. Equal number of cells were pooled from all mice in a group. 10,000 cells from each group were loaded on a 10X Genomics Next GEM chip and single-cell GEMs were generated on a 10X Chromium Controller. Subsequent steps to generate cDNA and sequencing libraries were performed following 10X Genomics’ protocol. Libraries were pooled and sequenced using Illumina NextSeq 200 and NovaSeq 6000 as per 10X sequencing recommendations. All samples had a sequencing yield of more than 237 million reads per sample.

The sequenced data were processed using Cell Ranger version 6.1.2 to demultiplex the libraries. The reads were aligned to *Mus musculus* mm10 and SCV2 (MN981442.1) genomes to generate count tables that were further analyzed and visualized using Seurat version 4.1.2 (*79*), dplyr (*80*), and ggplot2 (*81*), with minor corrections to made to the output display using ggpubr (*82*), and ggrepel (*83*). Differentially expressed genes were calculated using the Wilcoxon Rank Sum test with Bonferroni correction (*p*-value < 0.05, log fold change limit of 0.25 and the stipulation that genes must appear in at least 10% of cells in either cluster). Clusters were defined by manual expert classification, based upon cluster specific differentially expressed genes using the R Seurat::FindMarkers function. Total scRNAseq data were split into three groups (CD45neg, myeloid and lymphoid as per **FigS1A**) based on expert curation prior to downstream analyses and sub-clustering. Each subset had independent scaling, PCA, UMAP, neighbor, and cluster analysis performed with 30 PC, 30 dims, and resolutions of 0.4 and 0.6.

### Histology

Tissues were fixed in 10% neutral buffered formalin for 48-72 hours and embedded in paraffin. Embedded tissues were sectioned at 5µm and dried overnight at 42°C prior to staining. Specific anti-SCV2 immunoreactivity was detected using a SCV2 nucleoprotein antibody (Genscript) at a 1:1000 dilution. The secondary antibody was the Vector Laboratories ImPress VR anti-rabbit IgG polymer (cat# MP-6401). The tissues were then processed for immunohistochemistry using the Discovery Ultra automated stainer (Ventana Medical Systems) with a ChromoMap DAB kit (Roche Tissue Diagnostics cat#760–159). Detection of SARS-CoV-2 viral RNA was performed using the RNAscope 2.5 VS assay (Advanced Cell Diagnostic Inc.) on the Ventana Discovery ULTRA as previously described (*84*) and in accordance with the manufacturer’s instructions. Briefly, tissue sections were deparaffinized and pretreated with heat and protease before hybridization with the antisense probe RNAscope 2.5 VS prove-V-nCoV2019-S-sense (Advanced Cell Diagnostics Inc, cat 845709). Tissue slides were evaluated blindly by a board-certified veterinary pathologist for histopathological score and the presence/absence of SCV2 nucleoprotein in bronchiolar epithelial cells, pneumocytes and macrophages. Cell types were determined based on location and morphology.

### Statistical analyses

In all cases statistical analyses were performed on pooled data from 2-3 independent experiments, each with 3-8 mice per group. Details for each analysis performed are reported in the figure legends. All data points are shown on the graphs and no animals were excluded except due to technical failure as outline in the Study Design.

P-values were determined by Student’s unpaired *t*-test or Mann-Whitney test when comparing two groups, or by One-Way ANOVA with Tukey’s post-test or Kruskal-Wallis test with Dunn’s post-test when comparing three or more groups using GraphPad Prism software (v9). P-values below 0.05 were considered statistically significant.

### Figure visualization

Figures were generated in Adobe Illustrator and R (*85*) incorporating images from Biorender.com.

## Data availability

Single-cell RNA sequencing data generated in this study has been submitted to the NCBI GEO database and is available under the Accession ID GSE236601.

## Author contributions

Conceptualization: KLH, SN, CGF, AS. Methodology: KLH, SN, CSC, PJB. Investigation: KLH, SN, CSC, PJB, VP, EPA, SDO, DO, JL, MC. Resources: NLG, BAPL, RFJ. Data curation and analysis: KLH, SN, CSC, VP, SIO. Writing – original draft: KLH, AS. Writing – review and editing: KLH, SN, CSC, PJB, VP, CGF, DJ, OL, KDM-B, AS. Visualization: KLH, SN, CSC, SIO. Supervision: KDM-B, DJ, OL, AS.

Funding acquisition: AS.

## Acknowledgements

We are grateful to Dr. Daniel Barber (NIAID) and Dr. Sonja Best (NIAID) for discussion, Dr. Christine Nelson (NIAID) for assistance with setting up SCV2 inactivation protocols and Dr. David Eccles (Malaghan) for bioinformatics support. We also thank Virgilio Bundoc and Robert Thompson for technical assistance; the National Cancer Institute Genomics Core for single-cell RNA sequencing; Dr. Craig Martens and the RML Genomics Unit for viral sequencing; the NIAID Research Technologies Branch for assistance with flow cytometry and the NIAID animal care staff. KLH was partially supported by a Rutherford Postdoctoral Fellowship from Te Apārangi Aotearoa/Royal Society of New Zealand. This research was funded by the Intramural Program of NIAID, NIH.

## Supplementary figure legends

**Figure S1:**
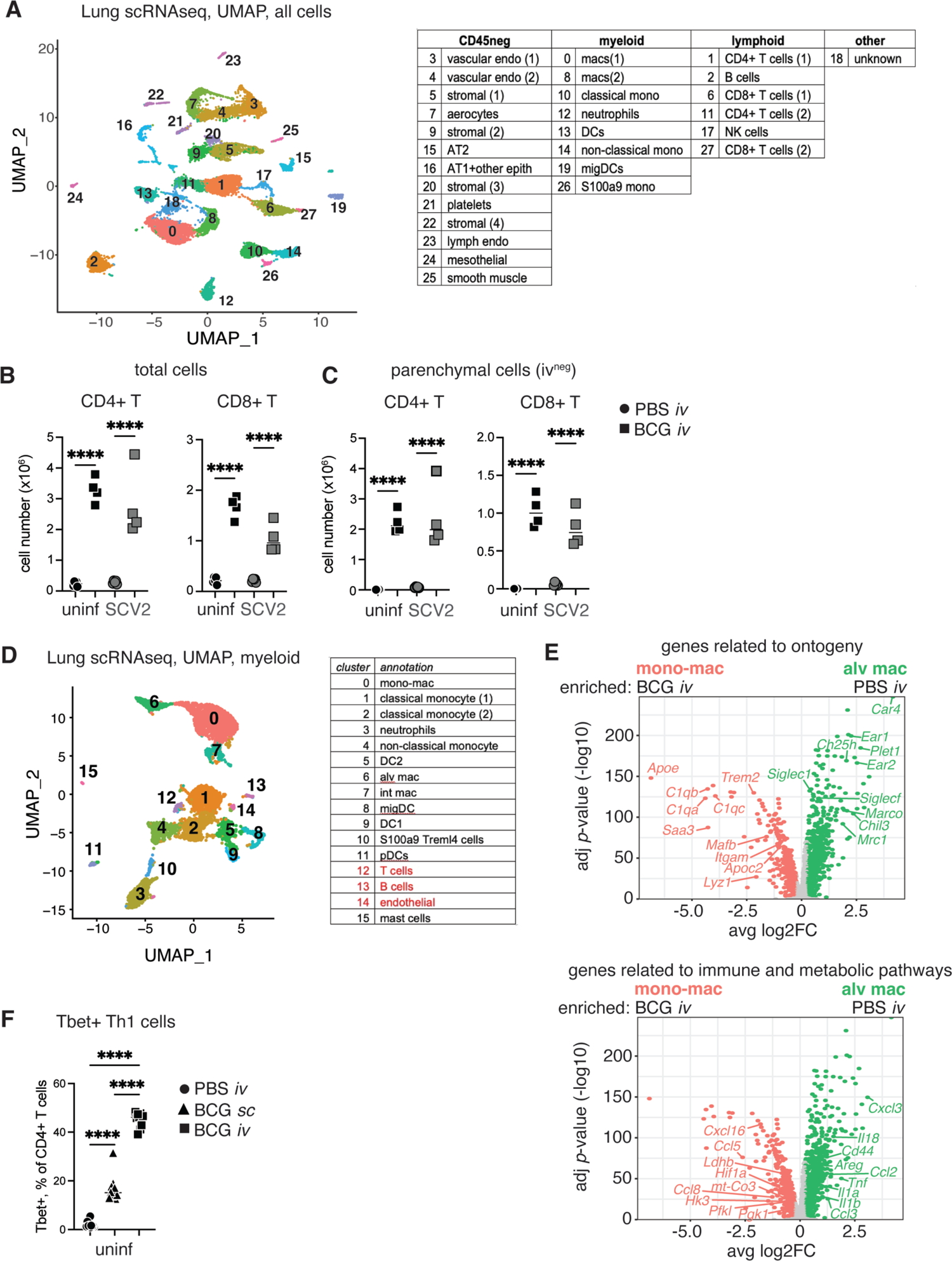
Th1 cells are enriched in the lung tissue following *iv* BCG. B6 mice were inoculated with BCG *sc* or *iv* 40-45 days prior to intranasal challenge with SCV2. Lungs were harvested 3 days after viral challenge. Control mice were not challenged with SCV2 (uninf). (**A**) UMAP representation of scRNAseq data of whole lung isolated from BCG and PBS animals challenged with SCV2 B.1.351. Clustering resolution: 0.6. Cell clusters were manually annotated and divided into 3 pools consisting of CD45-negative, myeloid or lymphoid linages (table on left). Cluster specific gene sets can be found in Supplementary Table 1. (**B**) Total number of lung CD4+ T cells and CD8+ T cells as determined by flow cytometry. (**C**) Number of CD4+ T cells and CD8+ T cells located in the lung parenchyma (panCD45 *iv* negative). Data are representative of 3 independent experiments with 4-5 mice/group. Statistical significance was determined by One-Way ANOVA with Tukey post-test. \*\*\*\**p*<0.0001. (**D**) UMAP representation of re-clustered myeloid cells for PBS or BCG treated animals. Clustering resolution: 0.4. Cell clusters were manually annotated (table on right). Cluster specific gene sets can be found in Supplementary Table 3. (**E**) Volcano plot shows DEGs between resident AM and monocyte-derived macrophage clusters with annotated cell ontogeny genes (upper panel) and genes related to immune and metabolic pathways (lower panel). Full gene lists and their associated FC and *p*-values can be found in Supplementary Table 4. (**F**) Number of lung Tbet+ CD4+ T cells as determined by flow cytometry. Data are pooled from 2 independent experiments with 4-5 mice/group. Statistical significance was determined by One-Way ANOVA with Tukey post-test. \*\*\*\**p*<0.0001.

**Figure S2:**
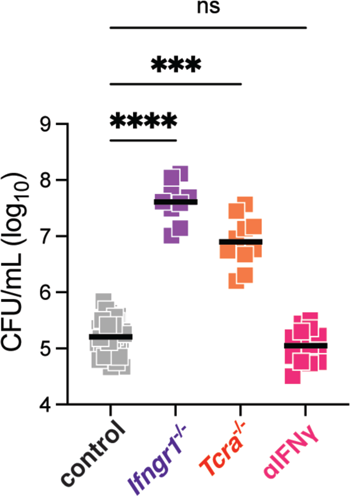
BCG CFU is not impacted by short-term IFNγ neutralization. Mice of the indicated genotypes were inoculated with BCG *iv* 40-45 days prior to intranasal challenge with SCV2. Lungs were harvested 3 days after viral challenge and plated on 7H11 agar to enumerate BCG CFU. Data are pooled from 2-3 independent experiments with 4-5 mice/group. Statistical significance was assessed by Kruskal-Wallis with Dunn’s post-test. Not significant (ns) *p*>0.05; \*\*\**p*<0.001; \*\*\*\**p*<0.0001.

**Figure S3:**
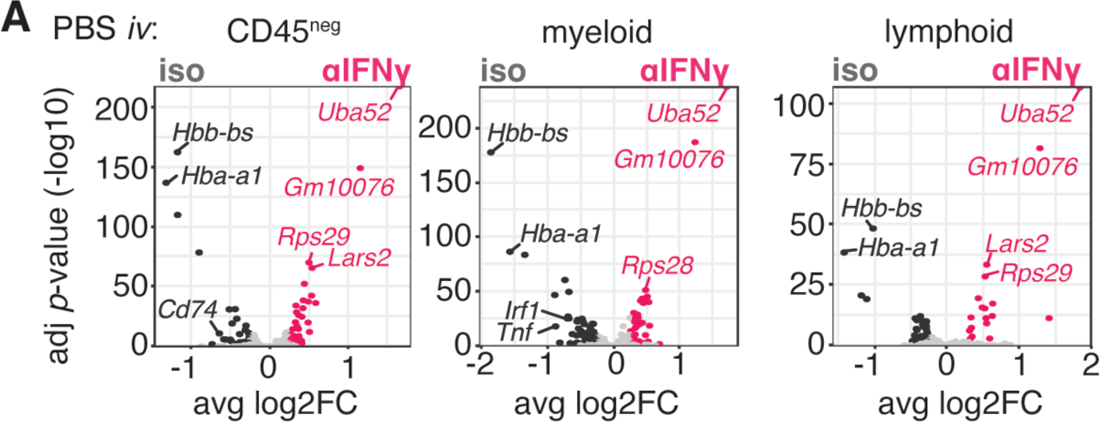
IFNγ neutralization minimally impacts gene expression 3 days post SCV2 infection. B6 mice were inoculated with PBS *iv* 40-45 days prior to intranasal challenge with SCV2 B.1.351. Infected animals received an IFNγ neutralizing antibody or isotype control 1 day prior to and 1 day following SCV2 instillation. Lungs were harvested 3 days after viral challenge. (**A**) Volcano plots show DEGs between isotype and anti-IFNγ treated mice inoculated *iv* with PBS across CD45^neg^, myeloid and lymphoid lineages that were manually annotated from the Seurat clustering showing in Fig 1C. Statistically significant DEGs are shown in dark gray or pink and were defined as FC>2.5 and *p*<0.05. Light grey points denote genes that did not reach statistical significance. Gene lists and their associated FC and *p*-values can be found in Supplementary Tables 8-10.

**Figure S4:**
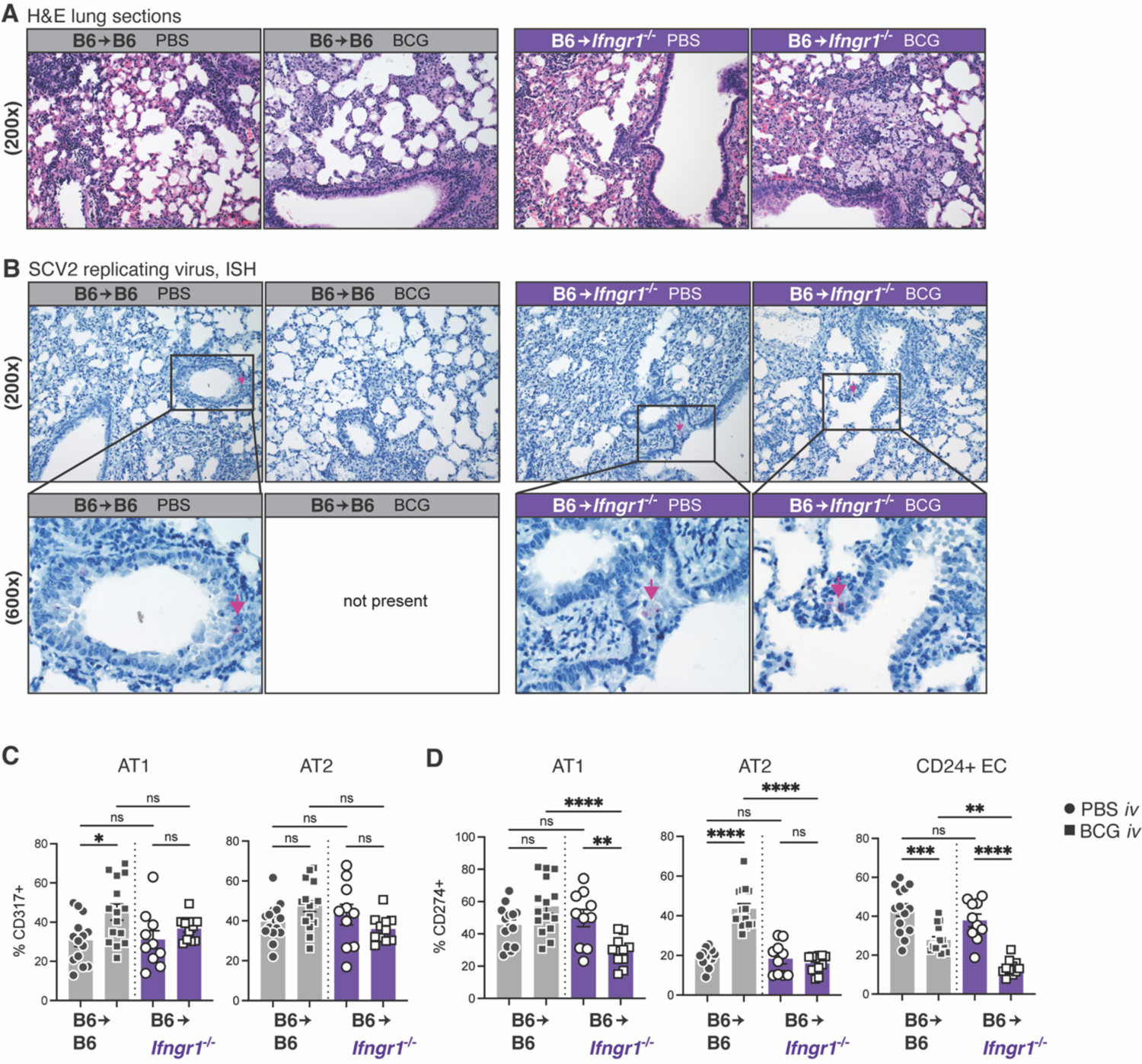
IFNγR1 signaling is required in the non-hematopoietic compartment for *iv* BCG induced protection against SCV2 infection. B6 congenic CD45.1+ mice were irradiated and reconstituted with either B6 CD45.2+ or *Ifngr1*-/- CD45.2+ bone marrow. Chimeras were inoculated with BCG or PBS *iv* 40-45 days prior to intranasal challenge with SCV2 B.1.351. Lungs were harvested 3 days after viral challenge. (**A**) Representative images of H&E-stained lung sections. (**B**) Example images of *in situ* hybridization of a probe targeting replicating SCV2. Data are representative of 2 independent experiments with 3-5 mice/group. (**C**) Expression of CD317 by AT1 and AT2 cells as determined by flow cytometry. (**D**) Expression of CD274 by AT1, AT2 and CD24+ epithelial cells as determined by flow cytometry. The epithelial cell gating strategy is shown in FigS5A. Data are pooled from 3 independent experiments with 3-4 mice/group. Statistical significance was assessed by One-Way ANOVA with Tukey post-test. Not significant (ns) *p*>0.05; **, *p*<0.01; ***, *p*<0.001; \*\*\*\**p*<0.0001.

**Figure S5:**
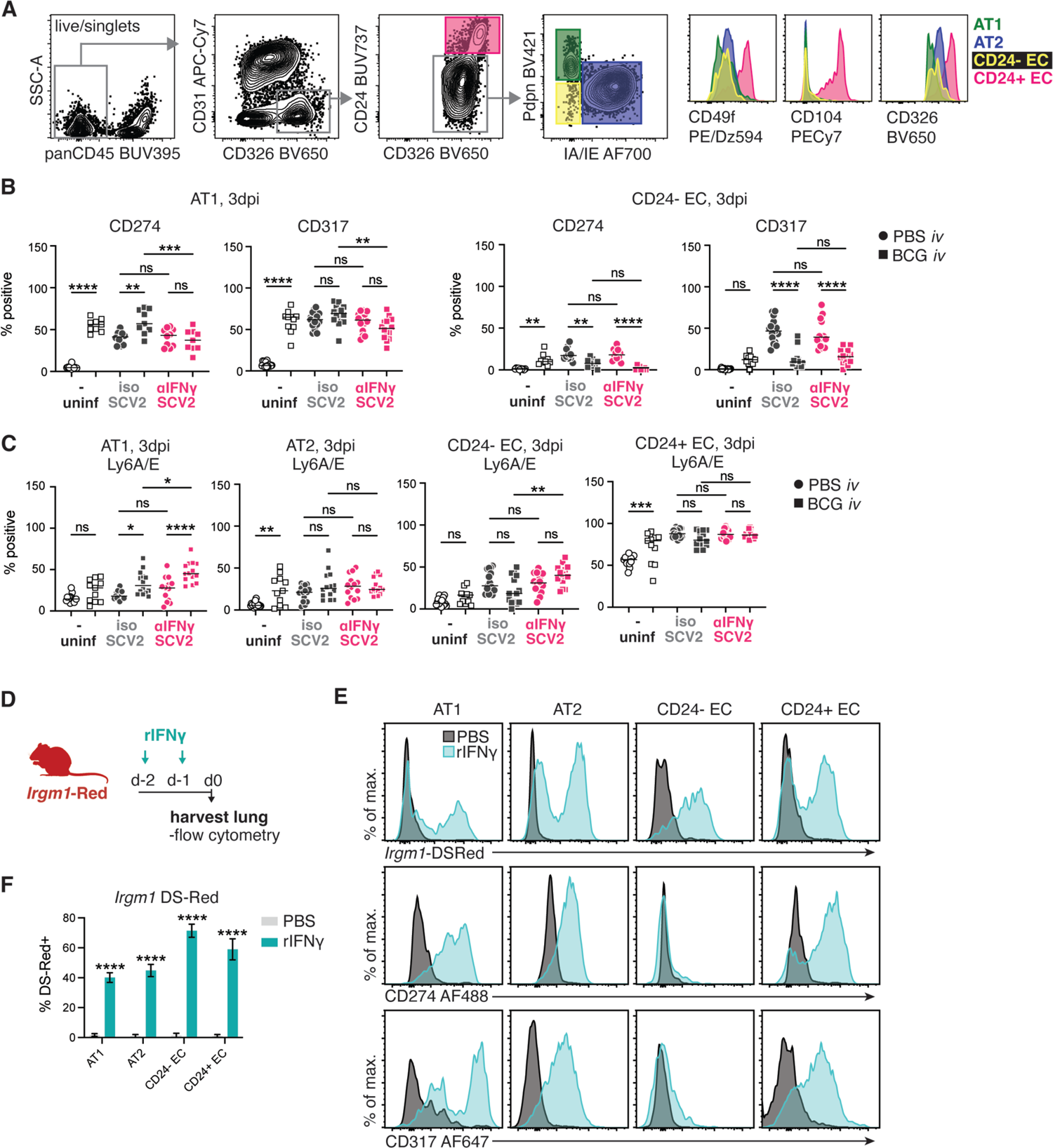
IFNγ induces expression of anti-viral markers in pneumocytes and CD24+ epithelial cells. (**A**) Gating strategy employed to identify pulmonary epithelial cell subsets. (**B-C**) B6 mice were inoculated with BCG or PBS *iv* 40-45 days prior to intranasal challenge with SCV2 B.1.351. Infected animals received an IFNγ neutralizing antibody or isotype control 1 day prior to and 1 day following SCV2 instillation. Lungs were harvested 3 days after viral challenge and the indicated epithelial cell types assessed for CD274, CD317 and Ly6A/E expression by flow cytometry. Data are pooled from 2-3 independent experiments with 4-5 mice/group. Statistical significance was assessed by One-Way ANOVA with Tukey post-test. Not significant (ns) *p*>0.05; \**p*<0.05; **, *p*<0.01; ***, *p*<0.001; \*\*\*\**p*<0.0001. (**D-F**) *Irgm1*-Red (M1-Red) mice were treated with PBS or rIFNγ intranasally on 2 consecutive days. Lungs were harvested 1 day after the last treatment. (**D**) Schematic of experimental protocol. (**E**) Representative expression profiles of *Irgm1* DS-Red, CD317 AF647 and CD274 AF488 across different epithelial subsets. (**F**) Expression of the *Irgm1* DS-Red reporter across different epithelial cell types. Data are pooled from 2 independent experiments with 3-5 mice/group. Statistical significance was assessed by unpaired t-test between PBS and rIFNγ treated B6 mice for each cell type. \*\*\*\**p*<0.0001.

**Figure S6:**
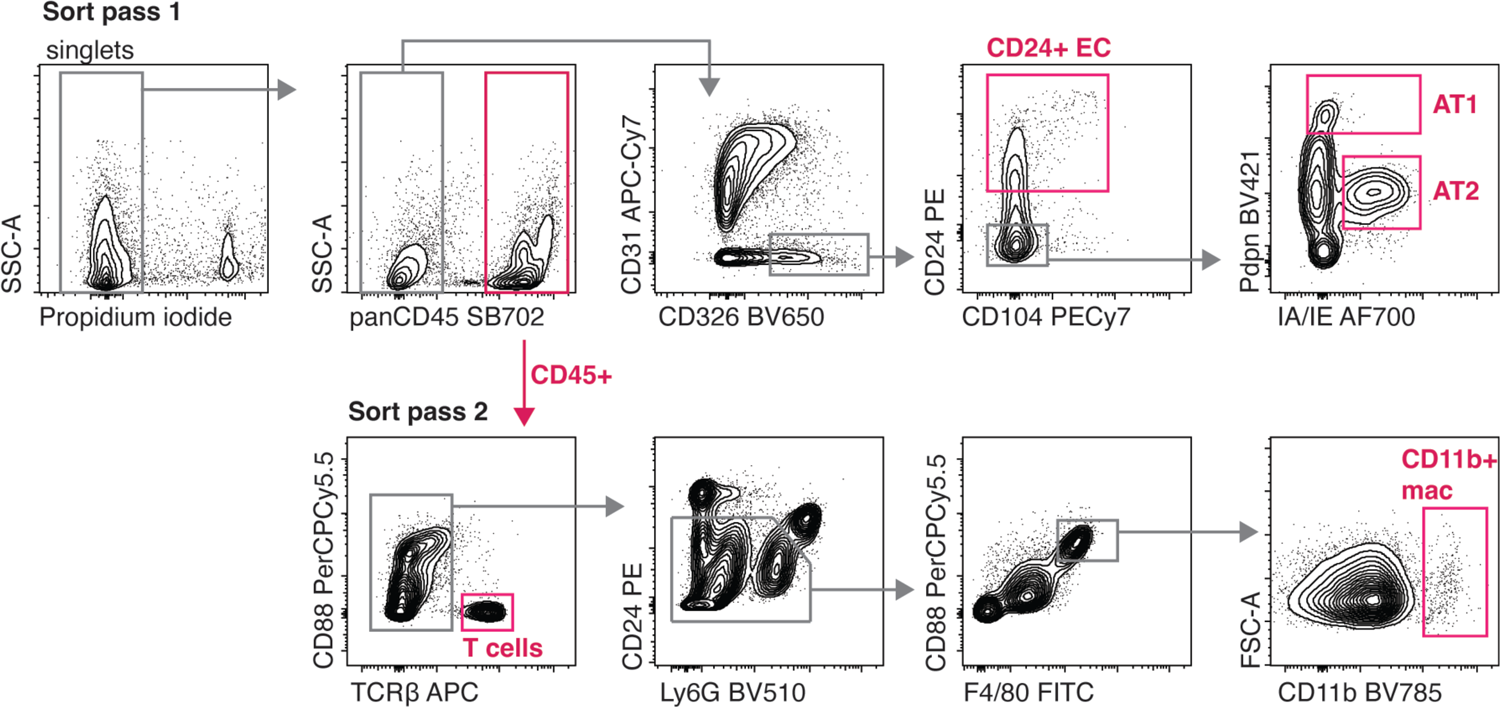
Cell sort gating strategy. Gating strategy employed to sort pulmonary epithelial cell subsets, T cells and CD11b+ macrophages from lung tissue of SCV2 infected K18-hACE2 mice.

## Notes

### Competing Interest Statement

The authors have declared no competing interest.

